# Cannabidiol confers neuroprotection against 6-OHDA toxicity by rescuing Nrf2 proteostasis and preserving mitochondrial integrity

**DOI:** 10.64898/2026.05.07.723512

**Authors:** Juan Jurado Coronel, Martin Duennwald

## Abstract

Oxidative stress and the progressive degeneration of dopaminergic neurons are key features of Parkinson’s disease (PD). The intrinsically disordered structure of the transcription factor Nuclear factor erythroid 2-related factor 2 (Nrf2), which coordinates the main cellular antioxidant response of the body, makes it highly susceptible to misfolding and aggregation under severe oxidative stress, compromising cellular survival. Cannabidiol (CBD) has potent neuroprotective properties, but its exact molecular mechanism within the dopaminergic redox environment remains unclear. In this study, we investigated the protective effects of CBD against 6-hydroxydopamine (6-OHDA)-induced toxicity in both undifferentiated and mature, post-mitotic differentiated SH-SY5Y cells. We found that CBD confers robust Nrf2-dependent neuroprotection against 6-OHDA. Importantly, we uncover a previously unexplored mechanism of neuroprotection by which CBD actively prevents the stress-induced sequestration of Nrf2 into insoluble cytoplasmic inclusions under oxidative stress. We find that CBD keeps Nrf2 in a soluble, functional state, increases Ser40 phosphorylation, restores nuclear localization, and drives the robust transcriptional upregulation of antioxidant enzymes. This targeted activation of Nrf2 effectively reduces intracellular ROS, significantly attenuates mitochondrial fragmentation, and decreases aberrant mitophagic activity. Overall, our results show that rather than merely scavenging reactive oxygen species, CBD directly increases Nrf2 activity during oxidative stress, enabling a sustained cytoprotective response. We thus identify CBD as a highly specific, targeted molecule with a high potential for neuroprotective therapy in PD.

## 1. Introduction

Parkinson’s disease (PD) is the second most common neurodegenerative disease, affecting 0.3% of the general population and almost 1% of people over age 60, presenting neuronal protein aggregates known as Lewy bodies (Poewe et al., 2017). Several pathomechanisms contribute to PD, one of the most important being oxidative damage to proteins and lipids, which causes mitochondrial dysfunction and further exacerbates oxidative stress (Hill *et al*. 2021; Burbulla *et al*. 2017). Despite the efficacy of dopamine-replacement therapies like Levodopa in managing symptoms, there remains no clinically approved agent capable of arresting or reversing this underlying oxidative degeneration in PD.

Cellular resilience against oxidative stress is primarily governed by the antioxidant pathway and the activation of its most important transcription factor, the nuclear factor erythroid 2-related factor 2 (Nrf2). This transcription factor maintains redox homeostasis in cells by regulating the expression of antioxidant, anti-inflammatory, and cytoprotective genes (Kahroba et al., 2021).

Under normal conditions, Nrf2 binds to the Kelch-like ECH-associated protein 1 (Keap1), which suppresses Nrf2 by marking it for degradation via the ubiquitin-protease system (UPS) (Kobayashi et al., 2004). Keap1 dimers bind Nrf2 at two affinity sites—low (DLG) and high (ETGE) (Jiménez-Villegas et al., 2021). The N-terminal region of Keap1 associates with Cullin-3 to form an E3 ubiquitin ligase complex that ubiquitinates Nrf2 (Goode et al., 2016). Furthermore, Keap1 contains a double glycine repeat (DGR), a C-terminal domain (CTR), an intervening region (IVR) with an NES, six Kelch repeats, and a Tramtrack-Bric-a-Brac (BTB) domain (Saha et al., 2022). The DGR and CTR form a β-propeller structure interacting with the Neh2 domain of Nrf2 (Saha et al., 2020). The IVR has cysteines Cys272 and Cys288, which are vital for repressing Nrf2 binding, whereas the BTB mediates Keap1 dimerization and contains Cys151 residue that may be important for stress sensing (Ogura et al., 2010). The Neh2 domain mediates Nrf2 binding to Keap1, and along with Neh1, and Neh1 contain transient structures that make Nrf2 a largely intrinsically disordered protein (IDP) (Karunatilleke et al., 2021). This disordered structure makes it susceptible to misfolding and inclusion formation (Ngo et al., 2022). Nrf2 misfolding and inclusion formation may thus be a mechanism by which excessive oxidative stress disables Nrf2, inhibiting the induction of the antioxidant response and exacerbating the cytotoxicity associated with oxidative stress (Ngo et al. 2022).

Upon exposure to electrophilic stress, Nrf2 dissociates from Keap1 -due to the oxidation of Cys151, Cys273, and Cys288 of Keap1-translocates to the nucleus, and binds to small MAF proteins (MafG, MafK, and MafF) and to Antioxidant Response Elements (ARE) to drive the expression of cytoprotective genes, such as *Heme Oxygenase-1 (HMOX1)* and *NAD(P)H Quinone Dehydrogenase 1 (NQO1)* (Kaur et al., 2024; Mahmoudi et al., 2019; Skibinski et al., 2017). However, in the context of PD, this adaptive response is often compromised as Nrf2-mediated expression of antioxidant genes is frequently downregulated in the Substantia Nigra pars compacta (SNpc) of PD patients (Kaur et al., 2024).

Cannabidiol (CBD), the major non-psychoactive phytocannabinoid from *Cannabis sativa*, has emerged as a promising neuroprotective candidate due to its potent antioxidant and anti-inflammatory properties (Atalay et al., 2022). Unlike Delta9-tetrahydrocannabinol (THC), CBD lacks intrinsic psychotropic activity and has a favourable safety profile (Campos et al. 2016).. This lack of psychoactive effect is due to its limited interaction with cannabinoid receptors. The therapeutic and neuroprotective effects of CBD arises from interactions with other neuronal receptors such as the transient receptor potential vanilloid 1 (TRPV1), GABA_A_, 5-hydroxytryptamine subtype 1A (5-HT1A) (serotoninergic receptor), peroxisome proliferator-activated receptor gamma (PPAR-γ), adenosine receptors A1 and A2, lipoxygenase (LOX) and cyclooxygenase type 2 (COX2), and G-coupled receptors such as GPR55, GPR18, GPR119, GPR3, GPR6, and GPR12 (Jîtcă et al. 2023). Moreover, by acting as a partial agonist at the high-affinity state of dopamine D2 receptors, CBD may provide symptomatic relief in PD by mimicking endogenous dopamine signalling (Seeman, 2016). Beyond its receptor-level interactions, pre-clinical evidence demonstrates that CBD acts as a highly specific modulator of the mesolimbic dopamine system. Studies by Laviolette and colleagues have shown that CBD ameliorates aberrant dopamine neuronal activity and alleviates neurobehavioral deficits, bypassing the molecular pathways responsible for the adverse side effects typically associated with traditional pharmacological interventions. Furthermore, they showed that CBD achieves these protective effects by directly regulating intracellular signaling networks, including the mTOR/p70S6 kinase and ERK1-2 pathways (Renard et al., 2016).

CBD has antioxidant, neuroprotective, anxiolytic, cardioprotective, and anti-inflammatory properties (Alexander, 2016; Fernández-Ruiz et al., 2011; Pereira et al., 2021). As oxidative stress is a key factor in neurodegenerative diseases, including PD, it is crucial to understand how CBD regulates the oxidative environment. While numerous studies have demonstrated that CBD can attenuate oxidative damage by activating the Nrf2 pathway (Love et al. 2023; Atalay et al. 2022; Li et al. 2022; Sun et al. 2017; Atalay Ekiner et al. 2022), the underlying molecular mechanisms remain incompletely understood. Therefore, further investigation into the mechanism by which CBD reduces oxidative stress is crucial to exploring its therapeutic potential.

In the present study, we investigated the neuroprotective efficacy of CBD in a cellular model of PD. Specifically, we employ 6-OHDA-induced toxicity in undifferentiated and differentiated SH-SY5Y cells, a model that closely resembles the post-mitotic phenotype of dopaminergic neurons. While the broad cytoprotective properties of Cannabidiol are widely acknowledged, our study establishes a fundamentally novel biophysical mechanism of neuroprotection. Taking into account the previous demonstration that severe oxidative stress drives Nrf2 into insoluble cytoplasmic inclusions that disable the antioxidant response (Ngo et al., 2022), we show here for the first time that CBD actively prevents this proteostatic failure, keeping Nrf2 in a soluble, transcriptionally competent state during dopaminergic oxidative injury. By preserving a soluble, transcriptionally competent Nrf2, CBD increases the expression of critical antioxidant enzymes (e.g., HMOX1, NQO1), halting pathological mitochondrial fragmentation and reducing mitophagic flux to near-baseline levels. Ultimately, these findings provide novel mechanistic insights into the therapeutic potential of CBD as an antioxidant response modulator in PD.

## 2. Materials and Methods

### 2.1 Reagents and Antibodies

Cannabidiol (CBD) and 6-hydroxydopamine (6-OHDA) were purchased from Sigma-Aldrich (St. Louis, MO, USA). Because methanol is highly toxic to cells, it was removed from the CBD solution using a vacuum concentrator (Thermo Fisher Scientific Savant SpeedVac). CBD was dissolved in Dimethyl sulfoxide (DMSO) to a stock concentration of 20 mM, and 6-OHDA was prepared fresh in sterile saline containing 0.02% ascorbic acid to prevent oxidation. The primary antibodies were: anti-Nrf2 (16396-1-AP, Proteintech), anti-p-Nrf2 (Ser40) (ab76026, Abcam), anti-HMOX1 (10701-1-AP, Proteintech), anti-NQO1 (11451-1-AP, Proteintech), anti-SOD2 (13141, Cell Signaling), anti-Tubulin (ab6160, Abcam), anti-GAPDH (TA802519, Origene), and anti-Histone H3 (NBP2-36468, Novus Biologicals). Fluorescent secondary antibodies (Alexa Fluor 488, 594, and 680) and Horseradish peroxidase (HRP)-conjugated secondary antibodies were obtained from Thermo Fisher Scientific (Waltham, MA, USA).

### 2. 2. Cell Culture and Treatments

Human neuroblastoma SH-SY5Y cells were obtained from ATCC (Manassas, VA, USA). Cells were routinely maintained in high-glucose Dulbecco’s Modified Eagle Medium (DMEM) (319-005 LL, Wisent Bioproducts, Saint-Jean-Baptiste, QC, Canada) supplemented with 10% Fetal Bovine Serum (FBS; Wisent Bioproducts) and 1% Penicillin-Streptomycin (Thermo Fisher Scientific). Cells were kept in a humidified atmosphere at 37°C with 5% CO2. For all experiments, cells were used at passage numbers lower than 25 to ensure phenotypic stability. For differentiation, cells were treated with 1% FBS DMEM with 10 µM retinoic acid (RA) for 7 days and 20 ng/mL Brain-derived Neurotrophic Factor (BDNF) for another 7 days before experiments. For oxidative stress experiments, the culture medium was switched to high-glucose DMEM without sodium pyruvate to prevent ROS scavenging mediated by pyruvate. To assess neuroprotection, cells were divided into four experimental groups: (1) Vehicle Control (DMSO < 0.1%), (2) 6-OHDA only, (3) CBD only, and (4) CBD pretreatment + 6-OHDA. Before treatment, the media was replaced with low-fetal bovine serum (FBS) (1%) media to prevent rapid oxidation of 6-OHDA and CBD binding to the FBS. The pretreatment group was incubated with Cannabidiol (CBD) 5 µM for 24 hours. Following pretreatment, 6-OHDA was added to the relevant groups at a final concentration of 50 µM to induce oxidative stress and mitochondrial damage.

### 2.3. Cell Viability and Metabolic Activity Assays

To comprehensively assess cell survival and metabolic health, two complementary assays were employed.

#### 2.3.1. ATP Quantification (CellTiter-Glo 2.0)

Primary assessment of cell viability was performed by measuring intracellular ATP levels using the CellTiter-Glo® 2.0 Assay (Promega, Madison, WI, USA). SH-SY5Y cells were seeded in white-walled 96-well plates at a density of 1 × 10^4^ cells/well, allowing them to attach and divide for 48 h before treatment. Following treatments, plates were equilibrated to room temperature for 30 min. An equal volume of CellTiter-Glo 2.0 reagent (100 µL) was added to the culture medium in each well. Plates were shaken for 2 min to induce cell lysis, then incubated at room temperature for 10 min to stabilize the signal. Luminescence was recorded using a microplate reader (Cytation 5 Cell Imaging Multimode Reader, Biotek).

#### 2.3.2. Dehydrogenase Activity (CCK-8)

To confirm the CellTiter-Glo 2.0 results, metabolic activity was assessed using the Cell Counting Kit-8 (CCK-8) (Enzo Life Sciences, Inc., Farmingdale, NY, USA). Cells were treated as described above in transparent 96-well plates. At the end of the treatment period, 10 µL of CCK-8 solution was added to each well, and plates were incubated for 1 h at 37°C. The absorbance of the water-soluble formazan dye was measured at 450 nm using a microplate reader (Cytation 5 Cell Imaging Multimode Reader, Biotek). Viability for both assays was expressed as a percentage of the vehicle-treated control.

### 2.4. Western Blotting

Cells were seeded in 6-well plates at a density of 3 x 10^6^ cells/well, treated, and then lysed in RIPA buffer (Thermo Fisher Scientific) supplemented with 1X halt protease & phosphatase inhibitor cocktail (Thermo Scientific, 78440) and with 10 mM N-Ethylmaleimide (NEM) to irreversibly alkylate free thiols and prevent post-lysis oxidation of cysteine residues. Protein concentration was determined using the BCA Protein Assay Kit (Thermo Fisher Scientific). Equal amounts of protein (10–30 µg) were separated by Sodium dodecyl sulphate–polyacrylamide gel electrophoresis (SDS-PAGE) and transferred onto Polyvinylidene difluoride (PVDF) membranes using the Trans-Blot® Turbo™ Transfer System (Bio-Rad, Hercules, CA, USA) according to the manufacturer’s instructions. Membranes were blocked with 5% non-fat milk in 1X Tris-buffered saline with Tween 20 (TBST) for 1 h and incubated with primary antibodies overnight at 4°C. After washing 3x with TBST, membranes were incubated with HRP-conjugated secondary antibodies for 1 h at room temperature or at 37°C when the primary antibody was from Proteintech. Bands were visualized using SuperSignal™ West Pico PLUS Chemiluminescent Substrate (Thermo Fisher Scientific) and imaged on a ChemiDoc™ MP imaging system (Bio-Rad). Densitometric analysis was performed using ImageJ software (NIH).

### 2.5. Nuclear and Cytoplasmic Fractionation

Subcellular fractionation was performed using the NE-PER™ Nuclear and Cytoplasmic Extraction Reagents (Thermo Fisher Scientific) according to the manufacturer’s protocol. We confirmed the purity of the nuclear fractions by immunoblotting for Histone H3 (nuclear marker).

### 2.6. Sedimentation Assay (Solubility Analysis)

To assess Nrf2 protein solubility, cells were lysed in RIPA buffer (Thermo Fisher Scientific) containing protease and phosphatase inhibitors. Lysates were centrifuged at 15,000 x *g* for 20 min at 4°C. The supernatant (soluble fraction) was collected, and the pellet (insoluble fraction) was washed once with lysis buffer and resuspended in 2% SDS buffer with sonication. Both fractions were analyzed by Western blotting.

### 2.7. Immunofluorescence Microscopy

Cells seeded on 15 mm glass coverslips were fixed after treatments with 2% paraformaldehyde for 15 min at 37°C, permeabilized with 0.2% Triton X-100, and blocked with 5% Normal Goat Serum (NGS). Cells were incubated with primary antibodies overnight at 4°C, followed by Alexa Fluor-conjugated secondary antibodies for 1 h at room temperature (RT). Nuclei were counterstained with Hoechst 33342. Imaging was performed using a Leica STELLARIS STED confocal microscope (Leica Microsystems, Mannheim, Germany) equipped with a White Light Laser (WLL) and Power HyD detectors, using an HC PL APO 63x or 100x/1.40 Oil CS2 objective. All images were acquired as Z-stacks with a step size of 0.8 µm spanning the full thickness of cells. Channels were acquired sequentially at 1024×1024-pixel resolution as 16-bit raw data files using Leica Application Suite X (LAS X) software, with identical laser power and gain settings maintained across all experimental groups.

### 2.8. Quantification of Nrf2 nuclear localization

Raw image files (.tiff format) were processed in batch using a custom ImageJ macro (Fiji distribution; (Schindelin et al., 2012)) implemented in the ImageJ macro language. For each acquisition, the two channels (Hoechst and Nrf2) were separated and saved as multi-page TIFFs, preserving the full Z-stack without any pixel-level modifications. Quantification of Nrf2 nuclear localization was performed using a per-slice analysis pipeline written as a custom ImageJ macro. For each Z-slice independently, the Hoechst channel was used to define the nuclear compartment as follows: the slice was smoothed with a Gaussian filter (σ = 2 px), background-subtracted using a rolling-ball algorithm (radius 100 px, sliding paraboloid), and thresholded using the Triangle method (Zack et al., 1977), and binarized. Holes within nuclei were filled, and connected components smaller than 3000 px were excluded as debris. The resulting binary mask defined the nuclear region for that slice.

Within each Z-slice, the Manders’ M2 colocalization coefficient (Manders et al., 1993) was calculated as the fraction of total Nrf2 fluorescence intensity that was located within the Hoechst-defined nuclear mask:

M2 = Σ I∼Nrf2∼(*x*,*y*) for (*x*,*y*) ∈ Nuclear mask / Σ I∼Nrf2∼(*x*,*y*) over the whole slice

Slices in which no nuclei passed segmentation (typically out-of-focus slices at the top or bottom of the stack) were excluded. The final M2 value reported for each image is the mean of the per-slice M2 values across all valid slices of that image. M2 values range from 0 (no Nrf2 in nuclei) to 1 (all Nrf2 in nuclei). A higher M2 indicates a greater fraction of cellular Nrf2 localized to the nucleus, interpreted as Nrf2 nuclear translocation, the hallmark of Nrf2 pathway activation (Itoh et al., 1999; Kobayashi et al., 2004).

### 2.9. Quantification of Nrf2 Inclusions (AggreCount)

To quantify Nrf2 inclusion formation, 16-bit raw images were acquired using a 100X objective and the Leica Application Suite X (LASX) software with identical laser power and gain settings maintained across all experimental groups. Automated image analysis was performed using the **AggreCount** macro for Fiji (ImageJ), as previously described (Klickstein et al., 2020). This workflow was selected for its ability to strictly segment high-intensity puncta from diffuse cytosolic background signal using a Difference of Gaussians (DoG) filter. These images were processed into two separate channels: Nuclei (Hoechst) and Aggregates (Nrf2, Alexa Fluor 488). Nuclei were segmented using automatic thresholding to define the cellular reference point. For Nrf2 inclusions, a Difference of Gaussians (DoG) filter was applied to the Alexa Fluor 488 channel to enhance small, high-frequency structures while suppressing low-frequency background noise. Resulting puncta were thresholded and filtered based on size (particles < 0.1 µm² were excluded as noise) to generate binary inclusion masks. Data were analyzed on a per-cell basis. For each experimental condition, at least 30 cells were analyzed across three independent replicates. The following metrics were calculated: Percentage of cells with inclusions: The fraction of the cell population containing at least one detectable Nrf2 inclusion; Inclusions per cell: The average number of discrete inclusions normalized to the total cell count; Inclusion area per cell: The total area of all segmented inclusions per cell (µm²), representing the total burden of sequestered protein.

### 2.10. Measurement of Intracellular ROS

Intracellular ROS levels were measured using the fluorescent probe CM-H2DCFDA (Thermo Fisher Scientific) in cells seeded in 96-well plates. As CM-H_2_DCFDA is a broad-spectrum oxidative stress indicator sensitive to multiple reactive species, including hydrogen peroxide, hydroxyl radicals, and peroxynitrite, the results reflect a general intracellular oxidative burden rather than the accumulation of any single ROS species. For measurement, cells were treated with 10 µM CM-H2DCFDA for 40 min before treatments, and fluorescence was measured at Ex/Em 485/535 nm. For flow cytometry measurements, cells were trypsinized, loaded with the probe, and analyzed on a BD FACSCelesta™ Cell Analyzer (BD Biosciences). Data were analyzed using FlowJo™ software (BD Biosciences).

### 2.11. Nrf2/ARE Reporter Assay

Transcriptional activity of Nrf2 was assessed using the ARE Reporter Kit - Nrf2 (Antioxidant Pathway) (Catalog #60514, BPS Bioscience, San Diego, CA, USA). Cells were seeded in 96-well plates and transiently transfected with the ARE luciferase reporter vector using Lipofectamine 3000 (Invitrogen), according to the manufacturer’s protocol. The kit contains a firefly luciferase gene under the control of ARE response elements, along with a constitutively expressed Renilla luciferase vector for normalization. Twenty-four hours post-transfection, cells were treated with CBD and 6-OHDA as described. After treatment, firefly and Renilla luciferase activities were measured sequentially using the Two-Step Luciferase (Firefly & Renilla) Assay System (Catalog #60683, BPS Bioscience) on a microplate reader (Cytation 5 Cell Imaging Multimode Reader, Biotek). Data were expressed as the ratio of Firefly to Renilla luminescence.

### 2.12. RNA Extraction and Quantitative Real-Time PCR (qPCR)

Total RNA was extracted using the PureLink™ RNA Mini Kit (Invitrogen) and reverse-transcribed into cDNA using the RevertAid H Minus First Strand cDNA Synthesis Kit (Thermo Fisher Scientific) according to the manufacturer’s protocol. qPCR was performed using SYBR Green Master Mix (Applied Biosystems, USA) on a QuantStudio™ 3 Real-Time PCR System (Applied Biosystems). Relative gene expression was calculated using the 2^−𝛥𝛥𝐶𝑡^ method, normalizing to *GAPDH* and *Ribosomal Protein Lateral Stalk Subunit P0* (*RPLP0*). Primer sequences are listed in **Supplementary Table 1**.

### 2.13. Mitochondrial Morphology and Mitophagy Analysis

After treatments, cells were stained with 100 nM MitoTracker™ Red CMXRos (Thermo Fisher Scientific) for 30 min at 37°C to label active mitochondria. Cells were then fixed, nuclei were stained with Hoechst 33342, and images were taken using a 63x objective. MitoTracker Red CMXRos was excited at 561 nm, and emission was collected at 570–620 nm. To ensure accurate morphological quantification, Z-stacks were acquired for each field of view with a 0.3 µm step size and subsequently compressed into 2D images using Maximum Intensity Projection in Fiji (ImageJ). Acquisition parameters (laser power, gain, and pinhole size) were kept constant across all experimental groups to allow for comparative analysis. Mitochondrial morphology parameters (branch length, form factor) were quantified using the “Mitochondria Analyzer” plugin in ImageJ. For each experimental condition, at least 30 cells were analyzed across three independent replicates. Analysis was performed on a per-cell basis to account for heterogeneity within the SH-SY5Y population.

For mitophagy analysis, cells were seeded into 12-well plates to achieve ∼60% confluency at the time of transfection. Cells were transiently transfected with the pLVX mCherry-Green Fluorescent Protein (GFP)-mtFIS1 plasmid (encoding the mCherry-GFP-Fis1 reporter) (Allen et al., 2013) using Lipofectamine 3000 according to the manufacturer’s instructions. Experiments were performed 48 hours post-transfection to allow adequate reporter expression. Cells were treated, fixed, and then imaged. To minimize spectral crosstalk, the reporter was imaged sequentially: GFP (ex: 488 nm/em: 500–550 nm) and mCherry (ex: 561 nm/em: 570–620 nm). All images were acquired as 16-bit raw data files using the Leica Application Suite X (LAS X) software platform. Identical laser power, gain, and offset settings were maintained across all experimental groups to ensure comparable quantification. Mitophagy was quantified by calculating the ratio of red-only puncta (mitolysosomes) to total mitochondrial area (yellow + red) per cell.

### 2.14. Statistical Analysis

Data are shown as mean ± standard deviation (SD) from at least three independent experiments. Statistical analysis was conducted using GraphPad Prism software (Version 9.5.1; GraphPad Software, San Diego, CA, USA). The normality of data distribution was evaluated with the Shapiro-Wilk test. Comparisons among multiple groups were made using one-way analysis of variance (ANOVA), followed by Tukey’s post hoc test for datasets with normal distributions. For datasets that did not follow a normal distribution, the Kruskal-Wallis test was used, followed by Dunn’s multiple comparison test. A p-value < 0.05 was considered statistically significant.

## 3. Results

### 3.1. Determination of optimal CBD concentration

We evaluated the basal cytotoxicity of cannabidiol (CBD) to identify a non-toxic dosing range. SH-SY5Y cells were treated with CBD concentrations ranging from 1 µM to 30 µM for 24 hours. CBD exhibited no significant cytotoxicity at lower concentrations (1 µM to 5 µM); however, significant reductions in cell viability were observed at doses of 7.5 µM (**Supplementary Figure 1**a). Based on our safety profiles, a maximum non-toxic dose of 5 µM CBD was selected to investigate its potential neuroprotective effects against oxidative stress.

### 3.2. CBD confers neuroprotection against distinct Parkinsonian toxins and mitigates 6-OHDA-induced toxicity in an Nrf2-dependent manner

We first determined whether CBD protects against three well-established neurotoxins used to model PD: 6-OHDA, a mitochondrial Complex I inhibitor that generates ROS directly through chemical auto-oxidation; Rotenone, a systemic Complex I inhibitor; and 1-methyl-4-phenylpyridinium (MPP+), the active metabolite of MPTP. Exposure to each toxin induced significant cytotoxicity in undifferentiated SH-SY5Y cells. Treatment with 50 µM 6-OHDA reduced cell viability to 67.82 ± 13.05% of the vehicle control (**Figure 1a**), while 10 µM Rotenone and 1 mM MPP+ decreased viability to 72.06%and 60.78%, respectively (**Figure 1b, 1c**). Pretreatment with 5 µM CBD demonstrated a robust, broad-spectrum protective effect. CBD reduced the 6-OHDA-induced cytotoxicity, preserving viability at 83.79 ± 11.45% of control levels (**Figure 1a**). Similarly, CBD effectively rescued cells from the toxic effects of mitochondrial inhibition induced by both Rotenone (87.51% of control levels; **Figure 1b**) and MPP+ (72.15% of control levels; **Figure 1c**).

**Figure 1.**
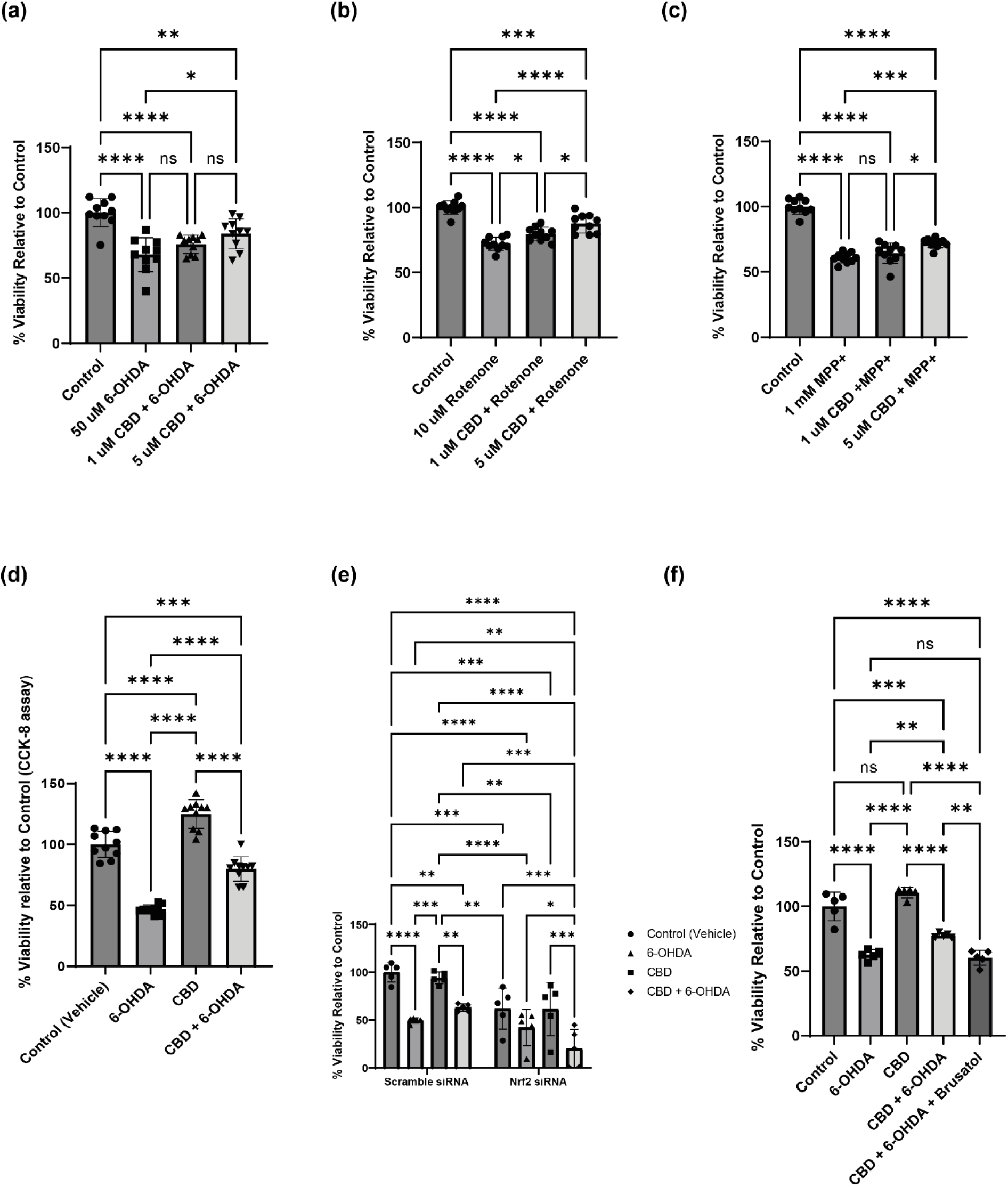
Cannabidiol (CBD) protects undifferentiated SH-SY5Y cells from 6-OHDA-induced toxicity in an Nrf2-dependent manner. Cell viability was assessed using the CellTiter-Glo® 2.0 luminescent assay and expressed as a percentage of the vehicle control, unless otherwise noted. Cells were pretreated with CBD for 24 h before co-treatment with 50 µM 6-OHDA for another 24 h. In (b) and (c), cells were pretreated with CBD for 24 h before co-treatment with 10 µM Rotenone (b) or 1 mM MPP+. **(a)** Dose-dependent protective effect of CBD (1 µM and 5 µM) against 6-OHDA toxicity in undifferentiated cells. **(b)** Protective effect of CBD (1 and 5 µM) against toxicity induced by 10 µM Rotenone. **(c)** Protective effect of CBD (5 µM) against toxicity induced by 1 mM MPP+. **(d)** Validation of viability using the CCK-8 colorimetric dehydrogenase activity assay. Cells were treated with 5 µM CBD alone, 6-OHDA alone, or the combination of both. **(e)** Genetic Nrf2 knockdown on CBD-mediated protection in undifferentiated cells. Cells were transfected with either Scramble non-targeting siRNA or Nrf2-specific siRNA before treatment with 5 µM CBD and 6-OHDA. **(f)** Pharmacological Nrf2 inhibition in undifferentiated cells. Cells were treated with 5 µM CBD and 6-OHDA in the presence or absence of the Nrf2 inhibitor Brusatol (200 nM). Data are presented as mean ± SD from *n*=10 (a-d) or *n*=5 (e-f) independent experiments performed in triplicate. Statistical significance was determined using one-way ANOVA followed by Tukey’s multiple comparisons test for (a), (b), (c), and (d), and two-way ANOVA followed by Tukey’s multiple comparisons test for (e) and (f). *p < 0.05, **p < 0.01, ***p < 0.001, ****p < 0.0001; ns, not significant. Comparisons indicate differences between specific treatment groups as shown by the brackets.

While CBD exhibited efficacy across all three PD-associated toxins, we selected 6-OHDA as the primary stressor for subsequent mechanistic studies. Unlike Rotenone and MPP+, which primarily target mitochondrial respiration, 6-OHDA mimics the specific pathology of dopaminergic degeneration by auto-oxidation, generating intracellular reactive oxygen species (ROS) and quinones (Xu et al., 2024). This direct oxidative mechanism makes it an ideal model for investigating the Nrf2-mediated antioxidant response, which is crucial for our study. Moreover, a preliminary assessment indicated that 5 µM CBD provided superior protection compared to 1 µM, and thus this concentration was selected for subsequent experiments.

To establish a robust *in vitro* model of oxidative stress-induced toxicity, we first performed dose-optimization studies in undifferentiated SH-SY5Y cells. Cells were exposed to increasing concentrations of 6-OHDA (5-100 µM) for 24 hours. Cell viability assays revealed a dose-dependent cytotoxicity, with 50 µM 6-OHDA reducing viability to approximately 50% (**Supplementary Figure 1b**). This concentration, representing the approximate half-maximal inhibitory concentration (IC50), was selected for all subsequent experiments, as it provides an optimal and reversible therapeutic window to evaluate the efficacy of candidate neuroprotective agents (Carvalho et al., 2013; Schober, 2004; Xu et al., 2024).

Cell viability measured with the CCK-8 assay in 6-OHDA-treated cells declined to 46.59 ± 3.74% of that in Control cells, confirming widespread impairment of cellular metabolic function (**Figure 1d**). Moreover, the CBD pretreatment protective effect was also seen in the CCK-8 assay, where metabolic activity was restored to 79.84 ± 10.14% (**Figure 1d**). Importantly, treatment with CBD alone did not exhibit cytotoxicity; in fact, it enhanced metabolic activity compared to vehicle controls.

To determine if this protection from 6-OHDA by CBD was causally linked to the Nrf2 pathway, we performed experiments in cells with reduced Nrf2 activity. First, cells were transfected with siRNA for Nrf2 knockdown. Successful depletion of Nrf2 was confirmed via Western blot, which demonstrated a strong (∼77%) reduction in Nrf2 protein levels compared to Scramble siRNA controls **(Supplementary Figure 2a, b).** Notably, cell viability assays revealed a significant reduction in the basal survival of untreated control cells transfected with Nrf2 siRNA compared to their Scramble-transfected counterparts. This indicates that the baseline activity of Nrf2 is key for maintaining physiological redox homeostasis and cellular survival in SH-SY5Y cells, even before exposure to 6-OHDA. Furthermore, in these Nrf2-depleted cells, the protective efficacy of CBD against 6-OHDA was markedly reduced compared to Scramble controls (**Figure 1e**). To corroborate this, we utilized Brusatol, a pharmacological Nrf2 inhibitor (Harder et al., 2017). Co-treatment with this molecule also abolished the CBD-mediated rescue, reducing viability to levels indistinguishable from those in the 6-OHDA-only group (**Figure 1f**).

### 3.3. CBD confers partial neuroprotection in differentiated SH-SY5Y cells

While undifferentiated SH-SY5Y cells are a robust screening tool, they are highly proliferative, have a cancer-like metabolism, and thus do not resemble neurons in some important respects. We thus evaluated whether the protective effects of CBD were maintained in differentiated SH-SY5Y cells, which exhibit neuron-like features, such as reduced glycolytic activity and cell cycle arrest (Forster et al., 2016). RA/BDNF differentiation of SH-SY5Y is widely used to model neuron-like maturation; increased Microtubule-associated protein 2 (MAP2)/Hexaribonucleotide Binding Protein-3 (NeuN)/Synaptophysin and reduced HKII/LDHA, with elevated Peroxisome proliferator-activated receptor-gamma coactivator-1alpha (PGC-1α), support a more oxidative, neuron-like state (Xicoy et al., 2017).

Following a 14-day differentiation protocol with RA and BDNF (Simões et al., 2021), we first validated the physiological shift in these cells. RT-qPCR analysis confirmed a significant and expected upregulation in mRNA expression of MAP2, NeuN, and Synaptophysin compared with undifferentiated cells (Forster et al., 2016; Simões et al., 2021; Xun et al., 2012) (**Supplementary Figure 3a**). Furthermore, differentiation induced a significant upregulation of *PGC-1α*, a master regulator of mitochondrial biogenesis, suggesting a shift toward increased oxidative phosphorylation and greater reliance on mitochondria (Forster et al., 2016). To confirm this metabolic shift away from the glycolytic “Warburg” phenotype typical of undifferentiated neuroblastoma cells, we evaluated key glycolytic enzymes by Western blot. Differentiated cells displayed a significant reduction in the protein levels of Hexokinase II (HK II) and Lactate Dehydrogenase A (LDHA) (**Supplementary Figure 3b–d**). Collectively, these transcriptional and metabolic alterations confirm the successful induction of a mature, post-mitotic neuronal phenotype heavily reliant on oxidative metabolism, thereby validating these differentiated cells as a physiologically relevant model for assessing neuronal neurotoxicity.

We next evaluated whether the protective effects of CBD were maintained in differentiated SH-SY5Y cells. Cell viability was assessed using the CellTiter-Glo 2.0 assay under the same treatment paradigm used for undifferentiated cells. Exposure of differentiated cells to 50 µM 6-OHDA resulted in high cytotoxicity, reducing cell viability to 17.46 ± 9.89% of the vehicle control (**Figure 2a**). Treatment with 5 µM CBD alone did not significantly alter basal cell viability, confirming that CBD is not toxic to these post-mitotic neuron-like cells. Importantly, pretreatment with CBD significantly reduced 6-OHDA-induced cell death, restoring cell viability to 41.39% ± 15.81% (**Figure 2a**). While this represents a partial rescue compared to the near-complete protection observed in the undifferentiated cells, the effect remains significant. This confirms the protective mechanisms of CBD function effectively within the distinct metabolic and energetic constraints of a physiologically relevant neuron-like model.

**Figure 2.**
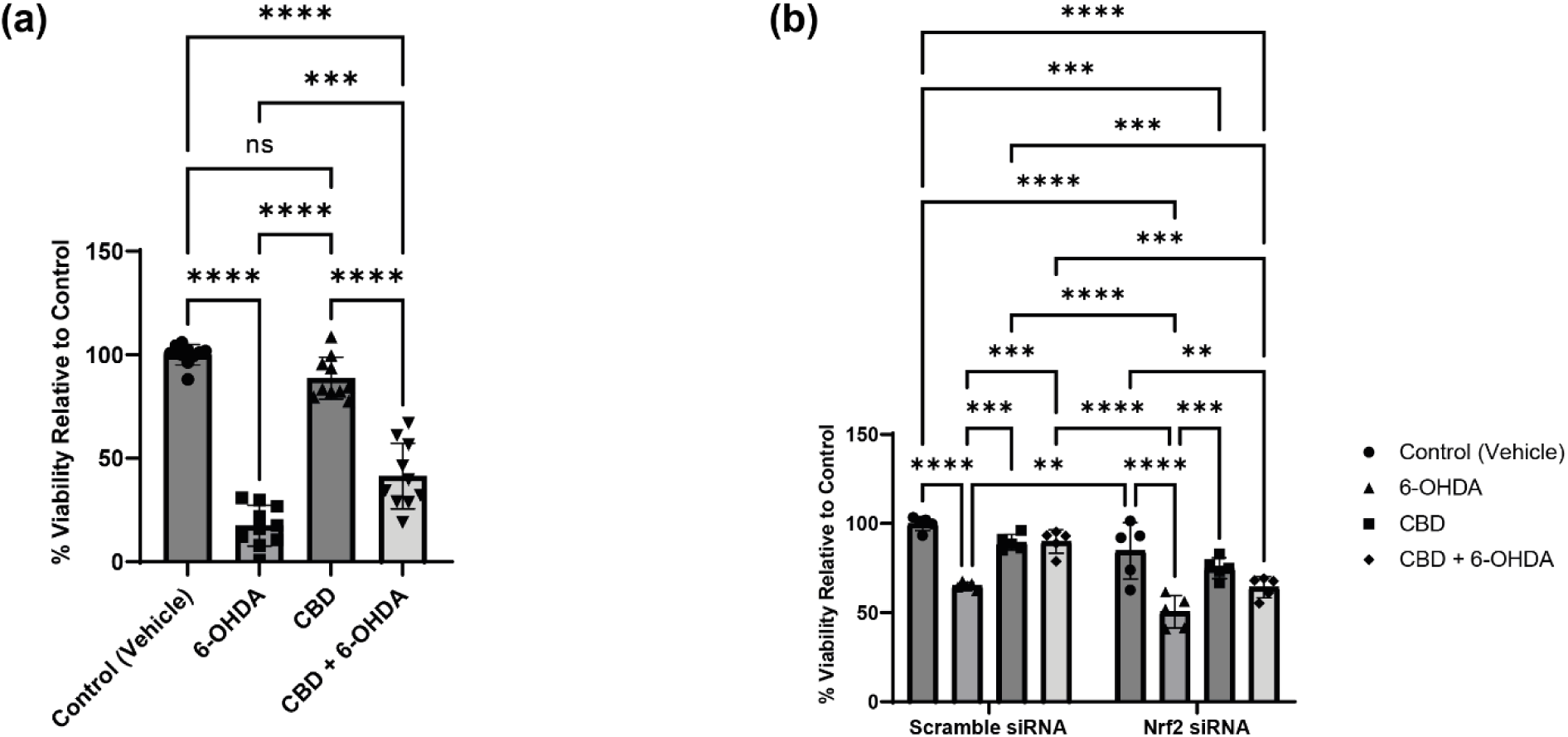
Neuroprotective efficacy of CBD is preserved in differentiated SH-SY5Y cells. SH-SY5Y cells were differentiated for 14 days (RA/BDNF) before treatment. Cell viability was determined using the CellTiter-Glo® 2.0 assay. **(a)** Protective effect of 5 µM CBD against 50 µM 6-OHDA toxicity in differentiated cultures. **(b)** Dependency on Nrf2 for neuroprotection in the differentiated state. Differentiated cells were transfected with Scramble or Nrf2-specific siRNA 48 h before treatment with CBD and 6-OHDA. Data are presented as mean ± SD from *n*=10 (a) or *n*=5 (b) independent experiments performed in triplicate. Statistical significance was determined using one-way ANOVA (a) or two-way ANOVA (b) followed by Tukey’s multiple comparisons test. *p < 0.05, **p < 0.01, ***p < 0.001, ****p < 0.0001; ns, not significant.

### 3.4. CBD reduces 6-OHDA-induced oxidative stress

Since Nrf2 activation primarily functions to neutralize ROS (Kaur et al., 2024) we determined whether the improvement in cell viability by CBD translated into a functional reduction in cellular oxidative stress. We first quantified global oxidative burden in undifferentiated SH-SY5Y cells. As expected, exposure to 6-OHDA led to a significant accumulation of intracellular ROS, resulting in a 1.47-fold increase in fluorescence intensity compared to controls (**Figure 3a**) (Schober, 2004). However, pretreatment with CBD significantly reduced ROS levels to levels statistically indistinguishable from those in the control (**Figure 3a**).

**Figure 3.**
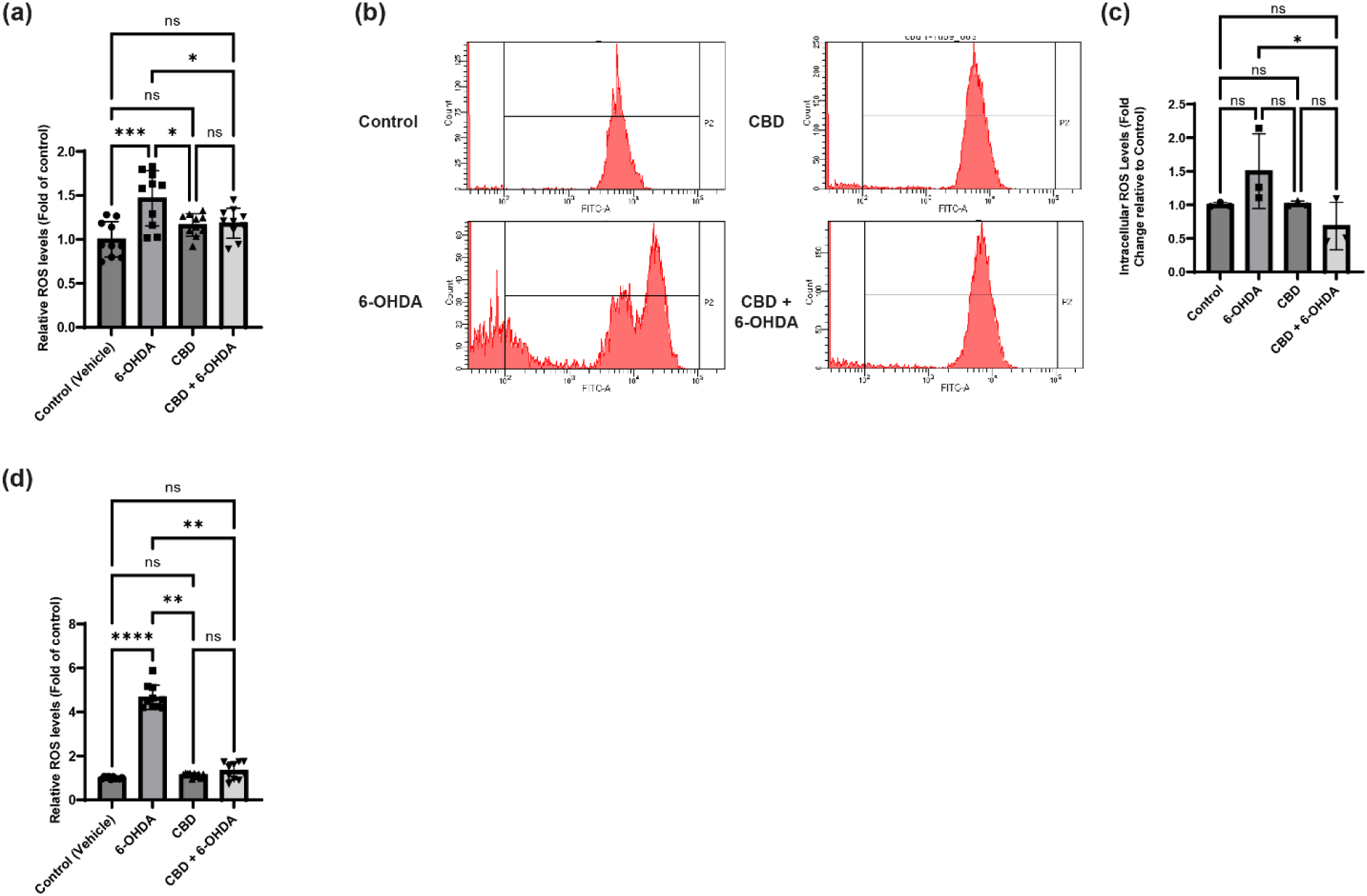
CBD attenuates 6-OHDA-induced oxidative stress. Unless otherwise noted, cells were pretreated with 5 µM CBD for 24 h before co-treatment with 50 µM 6-OHDA for another 24 h. **(a)** Intracellular ROS levels in undifferentiated SH-SY5Y cells were measured using the CM-H2DCFDA fluorescent probe in a 96-well plate format. Data are expressed as fold change relative to the vehicle control (*n*=10). **(b–c)** Flow cytometry analysis of intracellular ROS. **(b)** Representative histograms of CM-H2DCFDA fluorescence (FITC-A) in undifferentiated cells. **(c)** Quantification of the Mean Fluorescence Intensity (MFI), normalized to control (*n*=3). **(d)** Intracellular ROS levels in differentiated SH-SY5Y cells were measured using the CM-H2DCFDA fluorescent probe in a 96-well plate format. Data are expressed as fold change relative to the vehicle control (*n*=10). Data are presented as mean ± SD. Statistical significance was determined using one-way ANOVA followed by Tukey’s post-hoc test for (a, c) and the non-parametric Kruskal-Wallis test followed by Dunn’s multiple comparisons test for (d). *p < 0.05, **p < 0.01, ***p < 0.001, ****p < 0.0001; ns, not significant.

To validate this finding at single-cell resolution, we performed flow cytometry analysis using the same ROS detection probe (**Figure 3b**). Quantification of the mean fluorescence intensity (MFI) confirmed that CBD reduced intracellular ROS levels by approximately 54.5% compared to the 6-OHDA-only group, as the intracellular ROS levels in the 6-OHDA-only treatment were 1.5 ± 0.56 times the levels in the control group. These levels in the CBD pre-treatment were 0.68 ± 0.035 times those in the control group (**Figure 3c**). 6-OHDA treatment caused severe oxidative stress, whereas CBD pretreatment prevented this ROS accumulation, maintaining the population within the basal fluorescence range.

We next evaluated whether the antioxidant capacity of CBD was conserved in differentiated SH-SY5Y cells. Consistent with our observations in the undifferentiated model, exposure to 6-OHDA caused a massive intracellular oxidative burst, resulting in an almost 5-fold increase in relative ROS levels compared to vehicle controls (**Figure 3d**). Notably, this oxidative increase was substantially higher than the mild increase observed in undifferentiated cells, demonstrating a pronounced sensitivity of differentiated SH-SY5Y cells to 6-OHDA. Despite this severe oxidative stress, pretreatment with CBD decreased ROS accumulation. In the CBD + 6-OHDA group, relative ROS levels were only slightly higher than those in the control, indicating a significant reduction compared to the 6-OHDA-only condition. Furthermore, ROS levels in these CBD-rescued cells were statistically indistinguishable from both the vehicle control and CBD-alone groups, demonstrating that CBD effectively neutralizes 6-OHDA-induced oxidative stress.

### 3.5. CBD restores Nrf2 nuclear localization and phosphorylation

We next asked if CBD exerts its protective effects by activating the Nrf2 antioxidant defense pathway. To elucidate the mechanism underlying the observed neuroprotection, we examined the effect of CBD on the stability and subcellular distribution of Nrf2 **(Figure 4)**. We first assessed total Nrf2 protein levels in whole-cell lysates. Treatment with 6-OHDA alone did not significantly alter total Nrf2 levels compared with control, whereas pre-treatment with CBD resulted in a notable upregulation of Nrf2 steady-state levels (**Figure 4a, 4b**). We next analyzed Nrf2 phosphorylation at Serine 40, a modification essential for its stabilization and activation. 6-OHDA treatment increased p-Nrf2 levels by ∼27-fold. However, CBD pretreatment further potentiated this signal, significantly enhancing phosphorylation compared to 6-OHDA treatment alone (∼41-fold to control) (**Figure 4c, 4d**).

**Figure 4.**
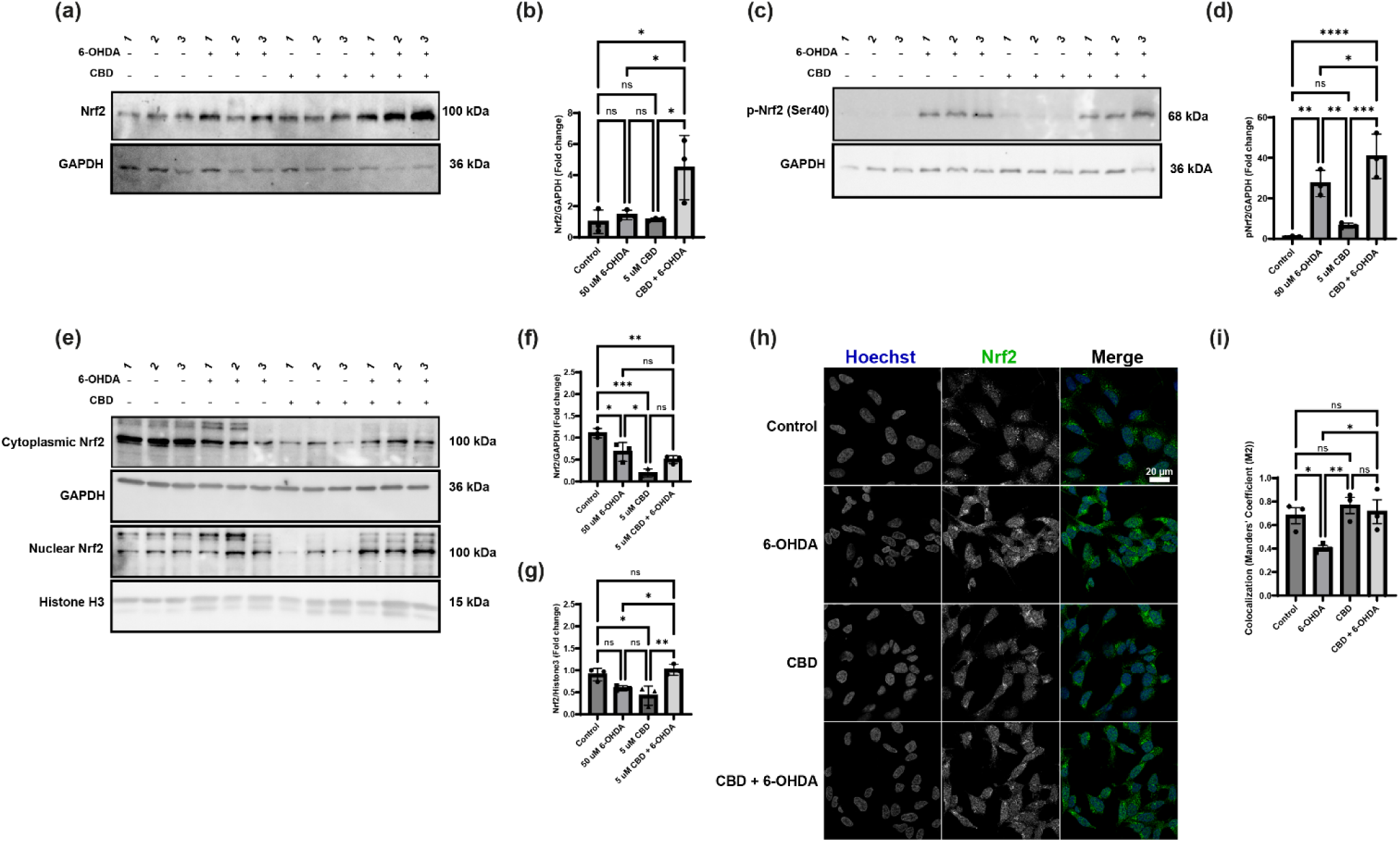
CBD promotes Nrf2 protein stabilization and nuclear translocation. Unless otherwise noted, data were obtained from undifferentiated SH-SY5Y cells treated with 5 µM CBD for 24 h. **(a–b)** Densitometric quantification of Nrf2 stabilization. Representative Western blots **(a)** and quantification **(b)** of total Nrf2 protein levels normalized to GAPDH. **(c–d)** Assessment of Nrf2 activation state. Representative Western blots **(c)** and quantification **(d)** of phosphorylated Nrf2 (p-Nrf2 Ser40) normalized to GAPDH. **(e–g)** Biochemical evidence of nuclear translocation via cellular fractionation. Representative Western blots **(e)** showing Nrf2 levels in cytoplasmic and nuclear fractions. GAPDH and Histone H3 were used as loading controls for the cytoplasmic and nuclear fractions, respectively. Densitometric quantification of Nrf2 in the cytoplasmic **(f)** and nuclear **(g)** fractions. **(h–i)** Visual confirmation of nuclear translocation via immunofluorescence. Representative confocal images **(h)** showing Hoechst (blue) and Nrf2 (green). Scale bar = 20 µm. **(i)** Quantification of Nrf2 nuclear translocation using Manders’ Colocalization Coefficient (M2). Data are presented as mean ± SD from *n*=3 independent experiments. Statistical significance was determined using one-way ANOVA followed by Tukey’s post-hoc test. *p < 0.05, **p < 0.01, ***p < 0.001, ****p < 0.0001; ns, not significant.

To confirm functional activation, we examined the subcellular distribution of Nrf2 by subcellular fractionation. In the cytoplasmic fraction, Nrf2 levels were significantly lower in all treatment groups than in the control (Figure 3e, 3f), suggesting either increased turnover, decreased expression, or transport to the nucleus (Kimura et al., 2007). Interestingly, 6-OHDA treatment led to a depletion of nuclear Nrf2, reducing levels to ∼50% of baseline (**Figure 4e, 4g**). This suggests that oxidative stress induced by 6-OHDA may compromise the nuclear translocation of Nrf2, possibly due to its aggregation in the cytoplasm (Ngo et al., 2022). Pretreatment with CBD prevented this loss, restoring nuclear Nrf2 levels to values indistinguishable from those in vehicle controls (**Figure 4e, 4g**).

Our biochemical findings were corroborated by immunofluorescence microscopy and quantitative colocalization analysis (**Figure 4h, 4i**). In control cells, Nrf2 exhibited a basal nuclear presence. Exposure to 6-OHDA significantly reduced the nuclear colocalization of Nrf2 (Manders’ Coefficient M2) compared to control (**Figure 4i**). However, in cells pretreated with CBD, this reduction was attenuated as the nuclear localization of Nrf2 was preserved at levels comparable to those in the control group (**Figure 4i**). Collectively, these data indicate that CBD protects cells not only by increasing Nrf2 levels and enhancing Nrf2 phosphorylation but also by preventing the stress-induced loss of Nrf2 from the nucleus.

### 3.6. CBD prevents the sequestration of Nrf2 into insoluble cytoplasmic inclusions

Our previous studies showed that Nrf2 forms cytoplasmic inclusions under oxidative stress (Ngo et al., 2022). Immunofluorescence analysis revealed that in cells treated with 6-OHDA, Nrf2 formed distinct, bright cytoplasmic inclusions (**Figure 5a**). Quantitative image analysis confirmed that 6-OHDA treatment significantly increased the presence of these inclusions compared to control cells. The percentage of cells containing inclusions rose significantly (**Figure 5b**), as did the number of inclusions per cell (**Figure 5c**) and the total inclusion area per cell (**Figure 5d**). Interestingly, pretreatment with CBD robustly reduced Nrf2 inclusion formation. In fact, in the CBD + 6-OHDA group, the Nrf2 inclusion pattern was similar to that in vehicle control, with inclusion formation returning to baseline levels (**Figure 5b–d**).

**Figure 5.**
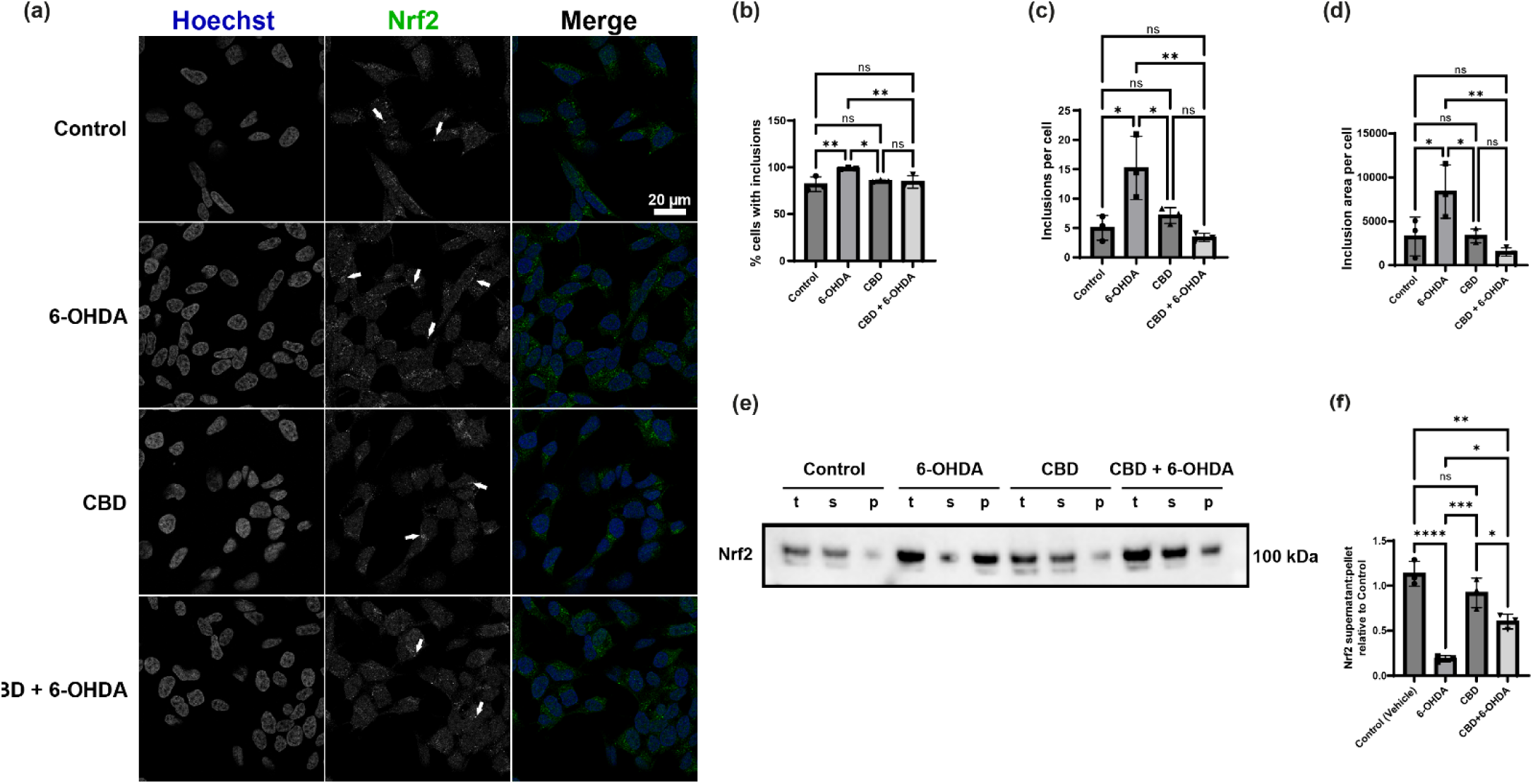
CBD prevents 6-OHDA-induced formation of insoluble Nrf2 inclusions. Undifferentiated SH-SY5Y cells were treated with vehicle, 50 µM 6-OHDA, 5 µM CBD, or a combination of both for 24 h, unless otherwise noted. **(a)** Representative confocal immunofluorescence images visualizing Nrf2 inclusions. Cells were stained with Hoechst (nuclei, blue) and Nrf2 (green). White arrows indicate dense Nrf2 inclusions, primarily observed in the 6-OHDA-treated condition. Scale bar = 20 µm. **(b–d)** Quantitative analysis of Nrf2 inclusions from confocal images. Graphs show **(b)** the percentage of cells containing visible inclusions, **(c)** the average number of inclusions per cell, and **(d)** the average total area of inclusions per cell. **(e)** Biochemical analysis of Nrf2 solubility using a sedimentation assay. Representative Western blot showing Nrf2 levels in total lysates (t), the soluble supernatant fraction (s), and the insoluble pellet fraction (p) obtained via high-speed centrifugation. **(f)** Densitometric quantification of the sedimentation assays. Data are expressed as the ratio of soluble (supernatant) Nrf2 to insoluble (pellet) Nrf2, relative to the vehicle control. Data are presented as mean ± SD from *n*=3 independent experiments. Statistical significance was determined using one-way ANOVA followed by Tukey’s post-hoc test. *p < 0.05, **p < 0.01, ***p < 0.001, ****p < 0.0001; ns, not significant.

To biochemically validate the nature of 6-OHDA-induced Nrf2 aggregation, we performed a sedimentation assay separating lysates into soluble (supernatant) and insoluble (pellet) fractions (**Figure 5e**). In vehicle-treated cells, Nrf2 was predominantly found in the soluble fraction, whereas 6-OHDA treatment caused a significant shift of Nrf2 into the insoluble fraction (**Figure 5f**), indicating Nrf2 aggregation. Pretreatment with CBD improved the solubility profile of Nrf2 (**Figure 5f**). Collectively, these data suggest that CBD reduces 6-OHDA-mediated Nrf2 aggregation, thereby enabling its nuclear translocation and, consequently, its function as a transcription factor.

### 3.7. CBD increases antioxidant gene and protein expression consistent with Nrf2 activation

We next verified whether CBD pretreatment indeed led to increased Nrf2-mediated transcription of antioxidant genes (Xu et al., 2024). We first used an ARE-driven luciferase reporter assay to measure promoter activity (Bakkar et al., 2015). While 6-OHDA treatment alone induced only a moderate increase in ARE activity (∼6.4-fold), pretreatment with CBD significantly potentiated this response, increasing luciferase activity to ∼11.7-fold relative to the control (**Figure 6a**).

**Figure 6.**
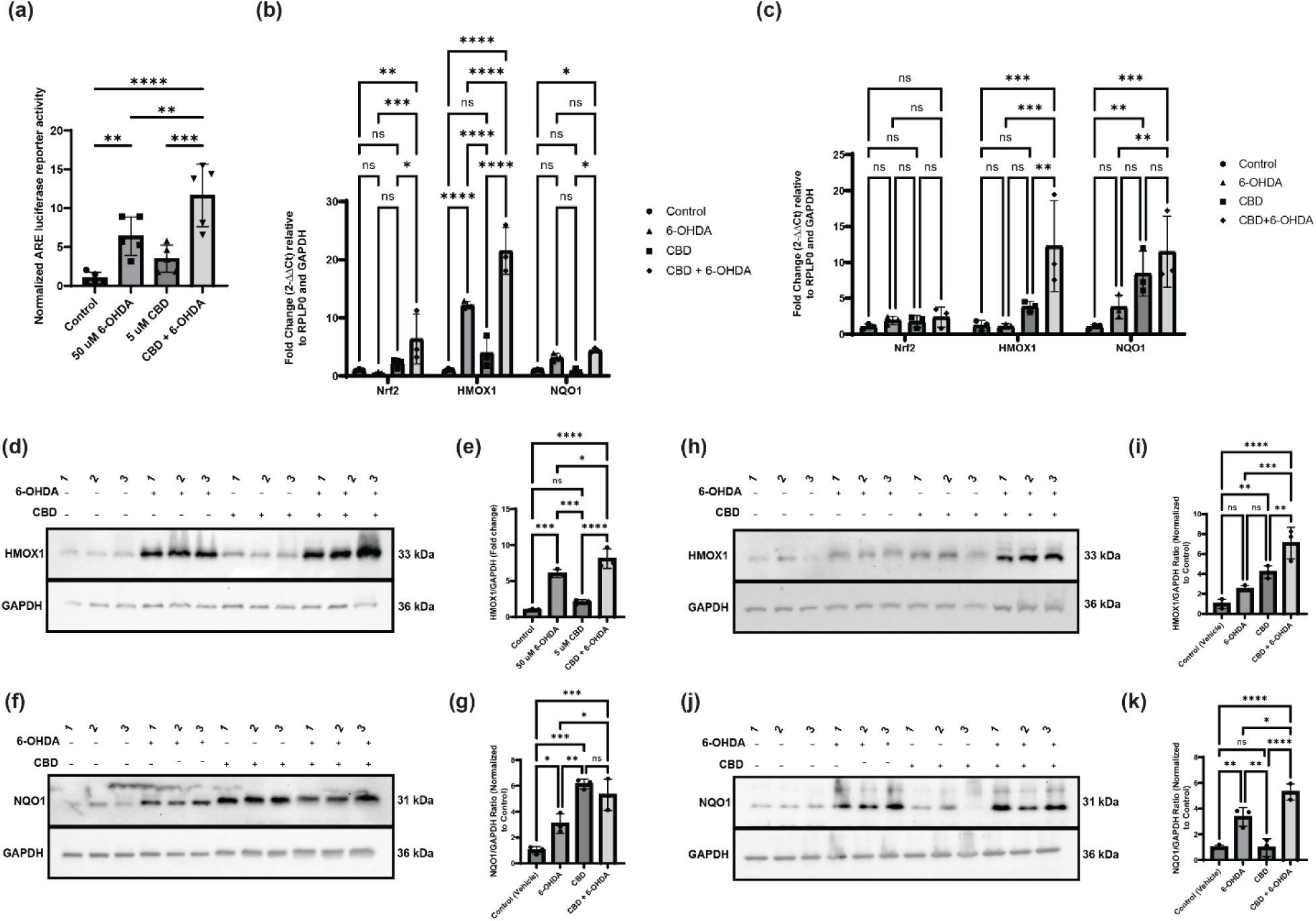
Nrf2 activation by CBD drives antioxidant gene transcription and protein expression. **(a)** CBD-induced Nrf2 transcriptional activity measured by an ARE-luciferase reporter assay. Data are expressed as normalized luciferase activity relative to vehicle control. **(b–c)** Relative mRNA expression of Nrf2-related antioxidant genes (*Nrf2*, *HMOX1*, *NQO1*, and *p62*) determined by RT-qPCR in undifferentiated **(b)** and differentiated **(c)** SH-SY5Y cells. Data are normalized to the housekeeping genes (*RPLP0* and *GAPDH*) and expressed as fold change (2^−𝛥𝛥𝐶𝑡^). **(d–g)** Western blot analysis of downstream target proteins in undifferentiated cells. Representative blots **(d)** and **(f)** and densitometric quantification **(e)** and **(g)** for HMOX1 and NQO1, respectively, normalized to GAPDH. **(h-k)** Western blot analysis of downstream target proteins in differentiated cells. Representative blots **(h)** and **(j)** and densitometric quantification **(i)** and **(k)** for HMOX1 and NQO1, respectively, normalized to GAPDH. Data are presented as mean ± SD from *n*=3 independent experiments. Statistical significance was determined using one-way ANOVA followed by Tukey’s post-hoc test for (a) and (e–k), and two-way ANOVA followed by Tukey’s post-hoc test for (b–c). *p < 0.05, **p < 0.01, ***p < 0.001, ****p < 0.0001; ns, not significant.

To confirm that this promoter activation translated to endogenous gene expression, we performed RT-qPCR experiments in both undifferentiated and differentiated SH-SY5Y cells. We selected three well-established Nrf2 targets—HMOX1, NQO1, and NFE2L2, because together they provide a comprehensive readout of the antioxidant response: HMOX1 is a highly sensitive indicator of global cytoprotective capacity, NQO1 specifically detoxifies the reactive quinones generated by 6-OHDA autoxidation, and NFE2L2, the gene encoding Nrf2, allows us to assess the transcriptional auto-regulation of the pathway (Johnson et al., 2008; Kaur et al., 2024). In undifferentiated cells, CBD pretreatment resulted in a robust upregulation of *HMOX1* (Heme Oxygenase-1), with mRNA levels increasing by ∼21.51-fold compared to the control and by **∼**9.33-fold compared to 6-OHDA alone (**Figure 6b**). Similarly, *NQO1* expression was significantly enhanced in the combination group compared to the control (**∼**4.38-fold), but not significantly higher than in the 6-OHDA-alone group (**Figure 6b**). Interestingly, *NFE2L2* (Nrf2) mRNA levels showed a more modest modulation, consistent with the premise that CBD primarily regulates Nrf2 at the level of protein stability rather than at transcriptional levels. While 6-OHDA treatment alone did not significantly alter *NFE2L2* mRNA levels compared to control, pretreatment with CBD followed by 6-OHDA resulted in a significant increase in expression compared to both the vehicle control and the 6-OHDA-only group. This indicates that CBD works synergistically with oxidative stress to drive *de novo* Nrf2 synthesis. We confirmed these transcriptional effects in differentiated cells, where CBD pretreatment significantly upregulated *HMOX1* (∼12.25-fold) and *NQO1* (∼11.47-fold) compared to the control, and ∼11.19- and ∼7.69-fold, respectively, compared to 6-OHDA only **(Figure 6c**).

We validated these Nrf2-mediated changes at the protein level by Western blotting, providing evidence that the observed transcriptional upregulation translates into the physical accumulation of stable, functional antioxidant enzymes capable of neutralizing ROS. In agreement with our mRNA analyses, CBD pretreatment significantly increased the steady-state protein abundance of HMOX1 (**Figure 6d, 6e**; ∼2.06-fold increase vs. 6-OHDA) and NQO1 (**Figure 6f, 6g**; ∼2.23-fold increase vs. 6-OHDA). This effect was maintained in differentiated cells, where CBD + 6-OHDA treatment resulted in a marked accumulation of HMOX1 (**Figure 6h, 6i**; ∼4.59-fold increase vs 6-OHDA) and NQO1 (**Figure 6j, 6k**; ∼1.6-fold increase vs 6-OHDA). Collectively, these data demonstrate that CBD not only stabilizes Nrf2 but also effectively drives the expression of critical downstream enzymes required to neutralize oxidative stress.

### 3.8. CBD preserves mitochondrial morphology

Oxidative stress compromises mitochondrial physiology, often shifting the balance towards fission and resulting in aberrant morphologies (Chan, 2006). We visualized mitochondrial architecture using MitoTracker Red CMXRos. This cell-permeant fluorophore accumulates in active mitochondria, dependent on their transmembrane potential, and covalently binds to mitochondrial proteins, enabling high-contrast structural resolution (Macho et al., 1996). Under basal conditions, undifferentiated SH-SY5Y cells displayed a reticular mitochondrial network characterized by long, tubular filaments (**Figure 7a**). By contrast, 6-OHDA treatment caused severe mitochondrial fragmentation, resulting in widespread swollen, irregular mitochondria, indicative of a catastrophic loss of network integrity, reflected by drastic morphological changes that typically precede excessive mitophagic clearance (Onishi et al., 2021; N. Sun et al., 2015). Quantitative morphological analysis confirmed this phenotype. 6-OHDA treatment significantly reduced the Mean Branch Length to 0.39 ± 0.07 µm, compared with 0.69 ± 0.04 µm in control cells (**Figure 7b**), confirming network fragmentation. Interestingly, 6-OHDA also significantly increased the Mean Form Factor (**Figure 7c**), where higher values indicate increasingly complex, irregular, or swollen shapes rather than the smooth, tubular structure of healthy mitochondria (H Koopman et al., 2005). Interestingly, CBD pretreatment preserved mitochondrial architecture in 6-OHDA-treated cells. Cells in the CBD + 6-OHDA group exhibited elongated, tubular mitochondria, similar to those in the control group. Quantification revealed that CBD significantly restored Mean Branch Length to 0.66 ± 0.16 µm (**Figure 7b**) and normalized the Form Factor (**Figure 7c**) to levels statistically indistinguishable from the vehicle control, indicating a remarkable rescue of mitochondrial structural integrity. Furthermore, in these undifferentiated cells, the total mitochondrial area did not differ significantly between the 6-OHDA and CBD + 6-OHDA treatment groups (**Supplementary Figure 4a**).

**Figure 7.**
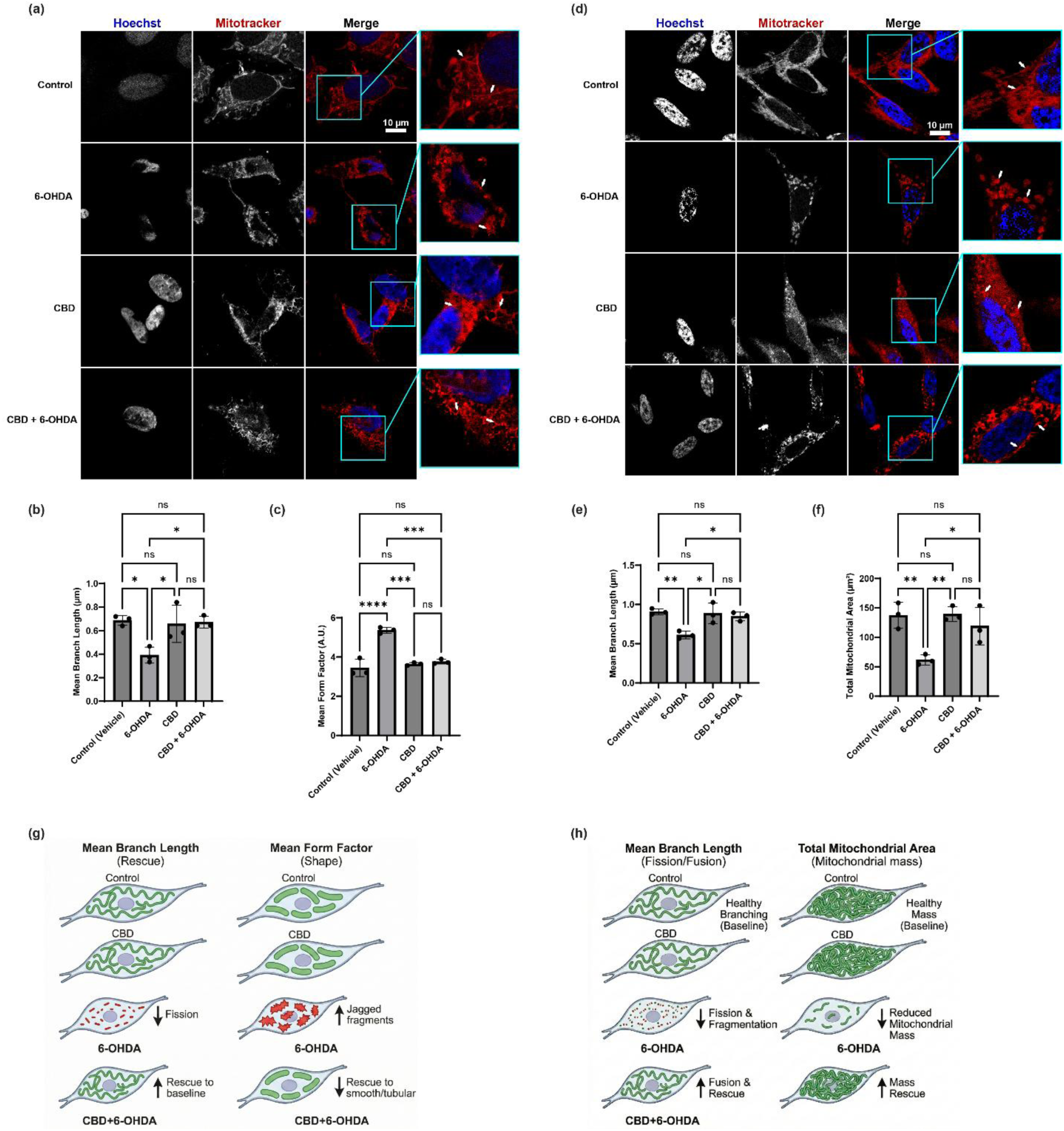
CBD attenuates 6-OHDA-induced mitochondrial fragmentation. Unless otherwise noted, cells were pretreated with 5 µM CBD for 24 h before co-treatment with 50 µM 6-OHDA for another 24 h. **(a)** Representative confocal microscopy images of mitochondrial morphology in undifferentiated SH-SY5Y cells visualized with MitoTracker™ Red CMXRos (red) and nuclei counterstained with Hoechst 33342 (blue). Scale bar = 10 µm. **(b–c)** Quantitative analysis of mitochondrial morphology parameters from confocal images: **(b)** Mean Branch Length, and **(c)** Mean Form Factor (indicating shape complexity). **(d)** Representative confocal microscopy images of mitochondrial morphology in differentiated SH-SY5Y cells. Scale bar = 10 µm. **(e–f)** Quantitative analysis of mitochondrial morphology parameters from confocal images, as shown in b: **(e)** Mean Branch Length, and **(f)** Total Mitochondrial Area. **(g-h)** Schematic summary illustrating the observed changes in mitochondrial morphology (fragmentation and shape changes) across the different treatment groups in undifferentiated **(g)** and differentiated **(h)** SH-SY5Y cells. For each experimental condition, at least 30 cells were analyzed across three independent replicates. Data are presented as mean ± SD. Statistical significance was determined using the non-parametric Kruskal-Wallis test followed by Dunn’s multiple comparisons test for (b-e) and one-way ANOVA followed by Tukey’s post-hoc test for (f). *p < 0.05, **p < 0.01, ****p < 0.0001; ns, not significant.

We then evaluated mitochondrial morphology and mass in differentiated SH-SY5Y cells **(Figure 7d).** Consistent with our previous observations, 6-OHDA exposure induced severe mitochondrial fragmentation in these differentiated cells, resulting in a significant reduction in Mean Branch Length to 0.61 ± 0.05 µm, compared with 0.91 ± 0.04 µm in control cells. Pretreatment with CBD effectively prevented this structural breakdown, maintaining Mean Branch Length at levels comparable to the baseline (0.85 ± 0.0.06 µm) (**Figure 7e**). Crucially, 6-OHDA treatment in differentiated cells also caused a severe depletion of mitochondrial mass, as evidenced by a significant decrease in Total Mitochondrial Area to 61.71 ± 8.88 µm2, compared with 137.4 ± 22.08 µm^2^ in control cells. CBD pretreatment significantly attenuated this loss of mitochondrial mass, successfully preserving Total Mitochondrial Area at 119.0 ± 31.83 µm2 (**Figure 7f**). Finally, we assessed shape complexity; however, unlike our findings in undifferentiated cells, the mean form factor did not differ significantly between the 6-OHDA and CBD + 6-OHDA conditions (**Supplementary Figure 4b**). Taken together, these data demonstrate that CBD provides robust, organelle-level protection by preventing the pathological fragmentation and mass reduction induced by 6-OHDA, thereby maintaining a stable, interconnected mitochondrial network across both undifferentiated and differentiated cells.

### 3.9. CBD normalizes mitophagic flux

Oxidative stress generates irreversibly damaged mitochondria that must be rapidly cleared to prevent cellular toxicity. We therefore investigated whether CBD-mediated protection also normalizes downstream mitochondria turnover. To quantify the clearance of damaged mitochondria by active mitophagy, we used the pLVX-mCherry-GFP-mtFIS1 reporter (Allen et al., 2013). This biosensor distinguishes between healthy mitochondria in the neutral cytosol (yellow: GFP+/mCherry+) and those delivered to the acidic lysosome for degradation by mitophagy (red: GFP-/mCherry+) (**Figure 8a**). In control cells, the mitochondrial network appeared predominantly yellow, indicating a stable population with low basal turnover. Treatment with 6-OHDA induced a significant increase in the appearance of red-only puncta, indicating mitolysosomes, reflecting the active clearance of damaged organelles (**Figure 8a**). Quantification of the Mitophagy Index (mitolysosomes per mitochondrial area) confirmed an increase in mitophagic flux in the 6-OHDA group compared to the control (**Figure 8b**). Conversely, in cells pretreated with CBD, the accumulation of mitolysosomes was significantly attenuated (**Figure 8b**). The mitochondrial network in the CBD + 6-OHDA group largely retained the yellow fluorescence signature of healthy organelles, with a mitophagy index comparable to baseline. This suggests that by preventing upstream oxidative damage and mitochondrial fragmentation, CBD obviates the need for excessive mitophagic clearance, preserving a healthy mitochondrial pool.

**Figure 8.**
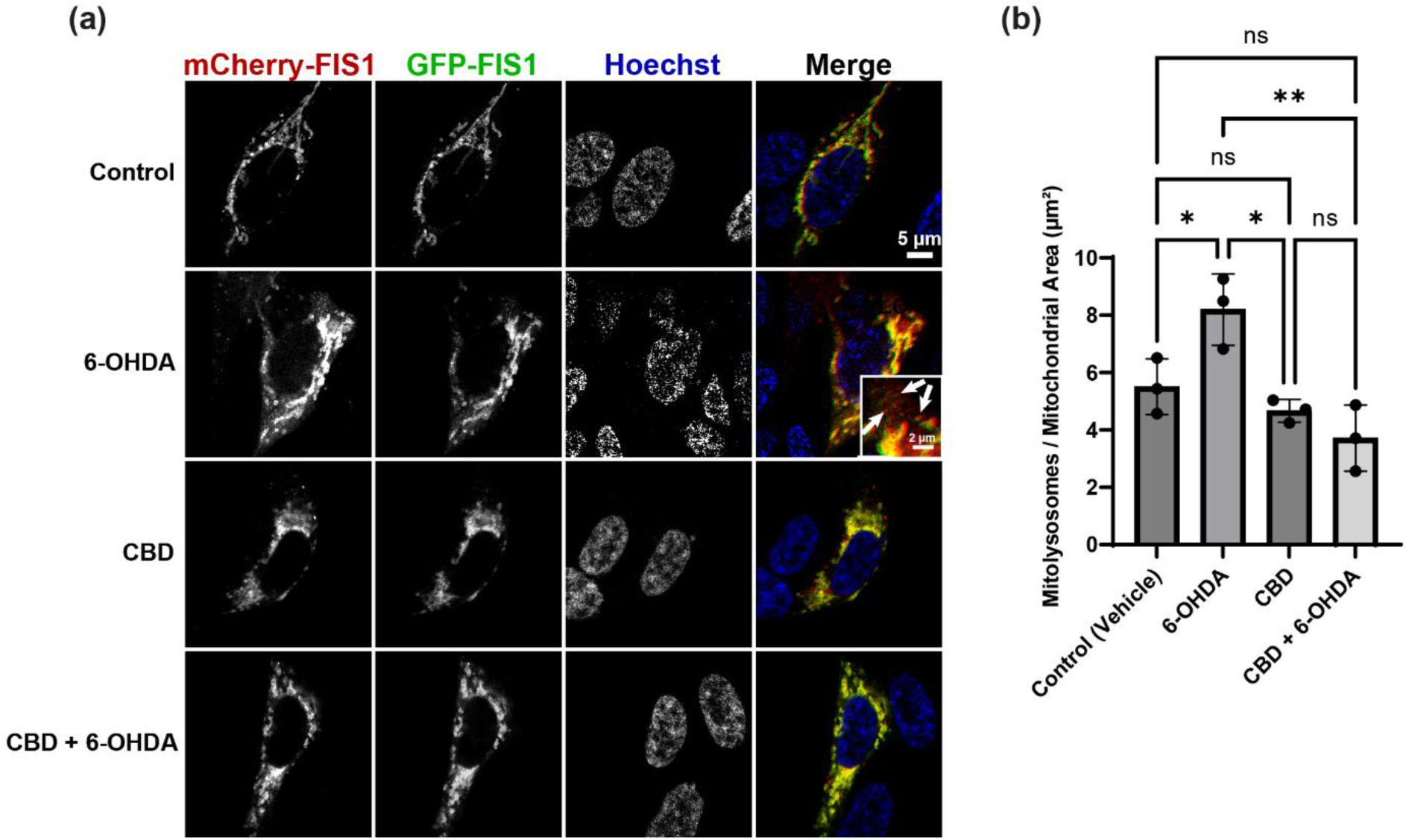
CBD attenuates 6-OHDA-induced mitophagy. **(a)** Representative confocal images of cells transfected with the tandem mCherry-GFP-Fis1 mitophagy reporter plasmid. Nuclei are stained with Hoechst (blue), mitochondria in the neutral cytosol appear yellow (green GFP + red mCherry), and mitochondria delivered to acidic lysosomes (mitolysosomes) appear red only (GFP quenched). The inset in the 6-OHDA panel highlights the accumulation of mCherry-only puncta (indicated by white arrows), representing active mitophagy events. **(b)** Quantification of the mitophagy index, calculated as the area of mitolysosomes (red-only puncta) divided by the total mitochondrial area per cell. The number of cells analyzed ranged from 17 to 36 across three independent replicates. Data are presented as mean ± SD. Statistical significance was determined using the non-parametric Kruskal-Wallis test followed by Dunn’s multiple comparisons test. *p < 0.05, **p < 0.01, ****p < 0.0001; ns, not significant.

To determine whether this localized mitochondrial protection was driven by compensatory upregulation of resident antioxidant defenses, we evaluated the protein expression of the primary mitochondrial superoxide scavenger, SOD2. Interestingly, western blot analysis revealed that total SOD2 expression remained relatively stable during neurotoxin exposure; 6-OHDA did not significantly alter SOD2 levels compared to vehicle controls. Furthermore, CBD pretreatment did not induce a significant upregulation of SOD2 compared to the 6-OHDA-only condition (**Supplementary Figure 5a**; ns). This indicates that the ability of CBD to rescue mitochondrial networks and halt pathological mitophagy is not dependent on de novo SOD2 synthesis but rather relies on upstream mechanisms, such as direct ROS neutralization or preservation of other critical Nrf2-dependent antioxidant systems.

## 4. Discussion

The progressive degeneration of dopaminergic neurons (DNs) in the SNpc is the central pathological feature of PD. While clinical interventions such as levodopa provide symptomatic relief, they fail to address the ongoing oxidative stress and the progressive collapse of endogenous antioxidant defenses that actively drive this nigrostriatal degeneration (McFarthing et al., 2024). Using SH-SY5Y cells treated with 6-OHDA, we demonstrated that CBD provides broad-spectrum neuroprotection. Of note, we find that CBD protects not merely through antioxidant scavenging but by maintaining Nrf2 in a soluble, nuclear, and transcriptionally competent state, thus enabling a coordinated cytoprotective gene expression program. We demonstrated that a pretreatment with CBD provides robust protection through the following mechanisms: (1) attenuating the accumulation of intracellular ROS, (2) increasing the phosphorylation of Nrf2 at Ser40 and restoring its nuclear localization, (3) preventing stress-induced sequestration of Nrf2 in insoluble cytoplasmic inclusions, (4) upregulating the transcriptional activation of the antioxidant enzymes HMOX1 and NQO1, and (5) preserving the integrity of the mitochondrial network while normalizing mitophagic flow. Importantly, our data go beyond the commonly reported observation that CBD can act as an antioxidant that scavenges ROS, as reported for many small molecule antioxidants (Olufunmilayo et al., 2023; Pisoschi & Pop, 2015; Xu et al., 2024) and instead begin to unravel a more specific mechanism by which CBD helps maintain Nrf2 in a soluble, functional state, enabling it to translocate to the nucleus in response to oxidative challenges. We propose that Nrf2 aggregation and impaired Nrf2 signalling are key contributors to the pathogenesis of PD, and CBD-mediated activation of Nrf2 might thus be a promising therapeutic intervention (Jakel et al., 2007; Robledinos-Antón et al., 2019; Yang et al., 2022).

We found that CBD provided significant protection against 6-OHDA, rotenone, and MPP+ in undifferentiated cells, consistent with previous results showing that its neuroprotective actions operate upstream of the specific toxin mechanisms (Fernández-Ruiz et al., 2013) and with its previously demonstrated antioxidant and anti-inflammatory profile (Alexander, 2016; Fernández-Ruiz et al., 2011). A 50 µM concentration of 6-OHDA reproduces the auto-oxidative ROS burst and mitochondrial dysfunction characteristic of dopaminergic degeneration in vitro (Blesa et al., 2012; Dias et al., 2013; N. Sun et al., 2015). The convergence of ATP strengthens the confidence in the protective effect of CBD- and dehydrogenase-based cell viability assays in our study, as these assays independently measure mitochondrial bioenergetics and cytosolic redox enzymes, respectively. It is important to note that protection was substantially reduced by both Nrf2-specific siRNA (∼77% knockdown) and the Nrf2 inhibitor Brusatol, establishing Nrf2 as a necessary factor for CBD-mediated survival. This positions CBD within a growing class of neuroprotective therapies that act through endogenous stress response systems rather than simple isolated scavenging.

The translational importance of this study is confirmed by the rescue in differentiated cells, which exhibit mature neuronal markers (MAP2, NeuN, Synaptophysin, PGC-1α) and reduced glycolytic enzymes (HKII, LDHA), reflecting increased mitochondrial dependence (Forster et al., 2016; Xun et al., 2012). Thus, the somewhat attenuated magnitude of the rescue compared to that found in undifferentiated cells could indicate a greater vulnerability to oxidative stress and mitochondrial dysfunction in cells with oxidative-dependent metabolism, rather than a loss of CBD activity per se, and is consistent with previous evidence that CBD modulates antioxidant responses in in vivo dopaminergic models and primary cortical neurons (García-Arencibia et al., 2007; Iuvone et al., 2004). The use of both undifferentiated and differentiated cells is clearly an experimental necessity, as the former allow for extensive mechanistic dissection under controlled conditions, and the latter provide a more physiologically relevant context in which Nrf2-dependent protective capacity remains intact, supporting the broader applicability of the findings of this study.

The Nrf2-dependent antioxidant response is the primary cellular defense against oxidative stress (Uruno & Yamamoto, 2023). Although 6-OHDA increased phosphorylation at Ser40, nuclear Nrf2 was paradoxically reduced, suggesting phosphorylation alone cannot trigger transcription when protein quality control is impaired. CBD pretreatment increased total Nrf2 and Ser40 phosphorylation more than 6-OHDA alone, and restored nuclear levels, possibly via Phosphoinositide 3-kinase / Protein kinase B (PI3K/Akt) and Mitogen-activated protein kinase / Extracellular signal-regulated kinase (MAPK/ERK) pathways through TRPV1 and GPR55, while also providing radical scavenging through phenolic hydroxyls (Atalay et al. 2020; Love et al. 2023). This proposed mechanism is strongly supported by the existing literature demonstrating the regulatory influence of CBD on central kinase networks. For instance, Laviolette et al. have shown that CBD functionally mitigates neurotoxic and psychotropic abnormalities by bidirectionally regulating ERK1-2 phosphorylation and directly modulating mTOR/p70S6 kinase signaling within dopaminergic pathways (Renard et al., 2016). Therefore, the robust CBD-mediated stabilization of Nrf2 proteostasis observed in our dopaminergic model is highly consistent with its ability to manipulate these exact downstream survival kinases. These actions reduce oxidative stress, facilitating nuclear translocation.

Functional activation of the Nrf2 pathway was indicated by increased ARE reporter activity and by HMOX1 and NQO1 expression at both mRNA and protein levels. HMOX1, a Nrf2 indicator that produces cytoprotective compounds, and NQO1, a detoxifying enzyme that metabolizes quinones, form an antioxidant defense relevant to the dopaminergic model, explaining the observed reduction in ROS. The modest NFE2L2 mRNA increase suggests CBD acts mainly post-translationally—stabilizing proteins, affecting localization and function—aligning with rapid Nrf2 regulation and strategies favouring stabilization over transcription, avoiding overexpression risks (Cuadrado et al., 2019; Kaur et al., 2024). NQO1 protein levels increase in undifferentiated cells pre-treated with CBD compared to those treated with 6-OHDA only, without higher mRNA levels, likely by reducing degradation or boosting translation, possibly via RNA-binding proteins like HuR (Park et al., 2011). Proteasome inhibition with MG-132 could confirm this, indicating that CBD influences NQO1 at the protein level and affects the entire Nrf2 pathway.

A central finding of this study is that oxidation-induced Nrf2 aggregation (Ngo et al., 2022) is recapitulated specifically in a dopaminergic oxidative environment modelled by 6-OHDA, and, importantly, that CBD actively prevents it. Nrf2 is an IDP, and its structural flexibility provides regulatory versatility but also makes it susceptible to misfolding and aggregation under constant oxidative stress (Karunatilleke et al., 2021; Ngo et al., 2022). Our findings indicate Nrf2 dysregulation in PD neurodegeneration, beyond transcriptional downregulation, to include a pathological deficiency in proteostasis, as assessed by protein oxidation, an aspect of PD pathophysiology that has received more attention. We propose that oxidative stress impairs the antioxidant defense system by sequestering Nrf2 in insoluble cytoplasmic inclusions. This produces a feedback of antioxidant failure that cannot be solved by phosphorylation alone. CBD pretreatment restored Nrf2 solubility and reduced all inclusion metrics to baseline.

CBD may maintain Nrf2 in a soluble, active state, through three potential pathways: first, the phenolic hydroxyl groups of CBD may directly scavenge reactive quinones and superoxide from 6-OHDA auto-oxidation (Atalay et al., 2020), reducing oxidative modifications on Nrf2 cysteine residues that trigger misfolding and inclusion formation (Karunatilleke et al., 2021; Ngo et al., 2022). Second, CBD might activate TRPV1 or GPR55 receptor-mediated signaling to trigger PI3K/Akt or MAPK/ERK cascades (Hind et al., 2016; Jîtcă et al., 2023), upregulating chaperones such as the Heat Shock Protein 70 (HSP70) to prevent Nrf2 misfolding. Selective pharmacological antagonism of these receptors would be expected to abolish the proteostatic effect without necessarily abolishing direct radical scavenging, providing the key experimental distinction between this pathway and the first. Third, CBD promotes autophagy via Sirtuin 1 (SIRT1)-dependent mechanisms (Kang et al., 2021; Pereira et al., 2021), which could decrease misfolded pools of Nrf2. Proteasome inhibition with MG-132 or autophagy blockade with bafilomycin A1 in the presence of CBD would distinguish protein quality-control–dependent from quality-control–independent routes. These models may act together, and understanding their relative roles remains a key focus of our research.

We also show that CBD preserved mitochondrial network integrity. CBD restored morphological parameters to control levels in undifferentiated cells and reduced mitochondrial mass loss in differentiated cells, which is consistent with findings in previous studies indicating the existence of CBD-mediated mitochondrial protection (Hind et al., 2016), and with a positive feedback model in which Nrf2-directed antioxidant gene expression stabilizes organelle homeostasis in cells dependent on oxidative phosphorylation (Dinkova-Kostova & Abramov, 2015; Forster et al., 2016; Xun et al., 2012). Moreover, we demonstrated that 6-OHDA increased mitophagic flux, consistent with PTEN-induced kinase 1 (PINK1)-Parkin-mediated compensatory clearing in acute PD models (Geisler et al. 2010; Kawajiri et al. 2010), and that CBD recovered the mitophagy index to control levels. We interpret this normalization as a consequence of decreased mitochondrial damage rather than a suppression of a protective function. CBD thus reduces the burden of dysfunctional mitochondria by preventing their oxidative fragmentation, which would otherwise require lysosomal clearance, thereby preserving a healthy, functionally competent mitochondrial pool.

To sum up, this study supports a model in which CBD concurrently enhances Nrf2 Ser40 phosphorylation, prevents stress-induced aggregation, restores nuclear translocation, directs HMOX1/NQO1 expression, reduces global ROS burden, preserves mitochondrial architecture, and normalizes mitophagic flux. This multilevel protection distinguishes CBD not only from simple ROS scavengers but also from single-target Nrf2 activators, such as sulforaphane or dimethyl fumarate, which modify Keap1 cysteines to increase Nrf2 activation but have no effect on Nrf2 proteostasis (Cuadrado et al., 2019; Satoh et al., 2022). The pleiotropic receptor profile of CBD, which includes 5-HT1A, PPAR-γ, TRPV1, and several G-protein-coupled receptors (GPCRs) (García-Gutiérrez et al., 2020), may involve various upstream nodes that converge on Nrf2 stabilization and proteostasis regulation (Campos et al., 2016; Jîtcă et al., 2023). This is advantageous in the context of a multifactorial disease such as PD, where single-mechanism therapies have consistently failed to improve the disease (Bloem et al., 2021; Poewe et al., 2017).

## Author Contributions

J.J.C. designed and performed research, analyzed data, generated figures, and wrote the paper. M.L.D. designed research, contributed reagents or analytical tools, and wrote the paper. All authors read and approved the submitted manuscript.

## Abbreviations Used

5-HT1A: 5-hydroxytryptamine subtype 1A
6-OHDA: 6-hydroxydopamine
ANOVA: analysis of variance
ARE: antioxidant response element
BDNF: brain-derived neurotrophic factor
BTB: Tramtrack-Bric-a-Brac
CBD: cannabidiol
CCK-8: Cell Counting Kit-8
COX2: cyclooxygenase type 2
CTR: C-terminal domain
DGR: double glycine repeat
DMEM: Dulbecco’s Modified Eagle Medium
DMSO: dimethyl sulfoxide
DNs: dopaminergic neurons
DoG: Difference of Gaussians
FBS: fetal bovine serum
GAPDH: glyceraldehyde-3-phosphate dehydrogenase
GFP: green fluorescent protein
GPCRs: G-protein-coupled receptors
HK II: hexokinase II
HMOX1: heme oxygenase-1
HRP: horseradish peroxidase
HSP70: heat shock protein 70
IC50: half-maximal inhibitory concentration
IDP: intrinsically disordered protein
IVR: intervening region
Keap1: Kelch-like ECH-associated protein 1
LAS X: Leica Application Suite X
LDHA: lactate dehydrogenase A
LOX: lipoxygenase
MAP2: microtubule-associated protein 2
MAPK/ERK: mitogen-activated protein kinase/extracellular signal-regulated kinase
MFI: mean fluorescence intensity
MPP+: 1-methyl-4-phenylpyridinium
MPTP: 1-methyl-4-phenyl-1,2,3,6-tetrahydropyridine
NEM: N-ethylmaleimide
NeuN: hexaribonucleotide binding protein-3
NGS: normal goat serum
NQO1: NAD(P)H quinone dehydrogenase 1
Nrf2: nuclear factor erythroid 2-related factor 2
PD: Parkinson’s disease
PGC-1α: peroxisome proliferator-activated receptor-gamma coactivator-1alpha
PI3K/Akt: phosphoinositide 3-kinase/protein kinase B
PINK1: PTEN-induced kinase 1
PPAR-γ: peroxisome proliferator-activated receptor gamma
PVDF: polyvinylidene difluoride
RA: retinoic acid
ROS: reactive oxygen species
RPLP0: ribosomal protein lateral stalk subunit P0
RT: room temperature
SD: standard deviation
SDS-PAGE: sodium dodecyl sulfate–polyacrylamide gel electrophoresis
SIRT1: sirtuin 1
SNpc: substantia nigra pars compacta
TBST: Tris-buffered saline with Tween 20
THC: Delta9-tetrahydrocannabinol
TRPV1: transient receptor potential vanilloid 1
UPS: ubiquitin-protease system
WLL: White Light Laser.

