## Supplemental Information for "Cannabidiol confers neuroprotection against 6-OHDA toxicity by rescuing Nrf2 proteostasis and preserving mitochondrial integrity"

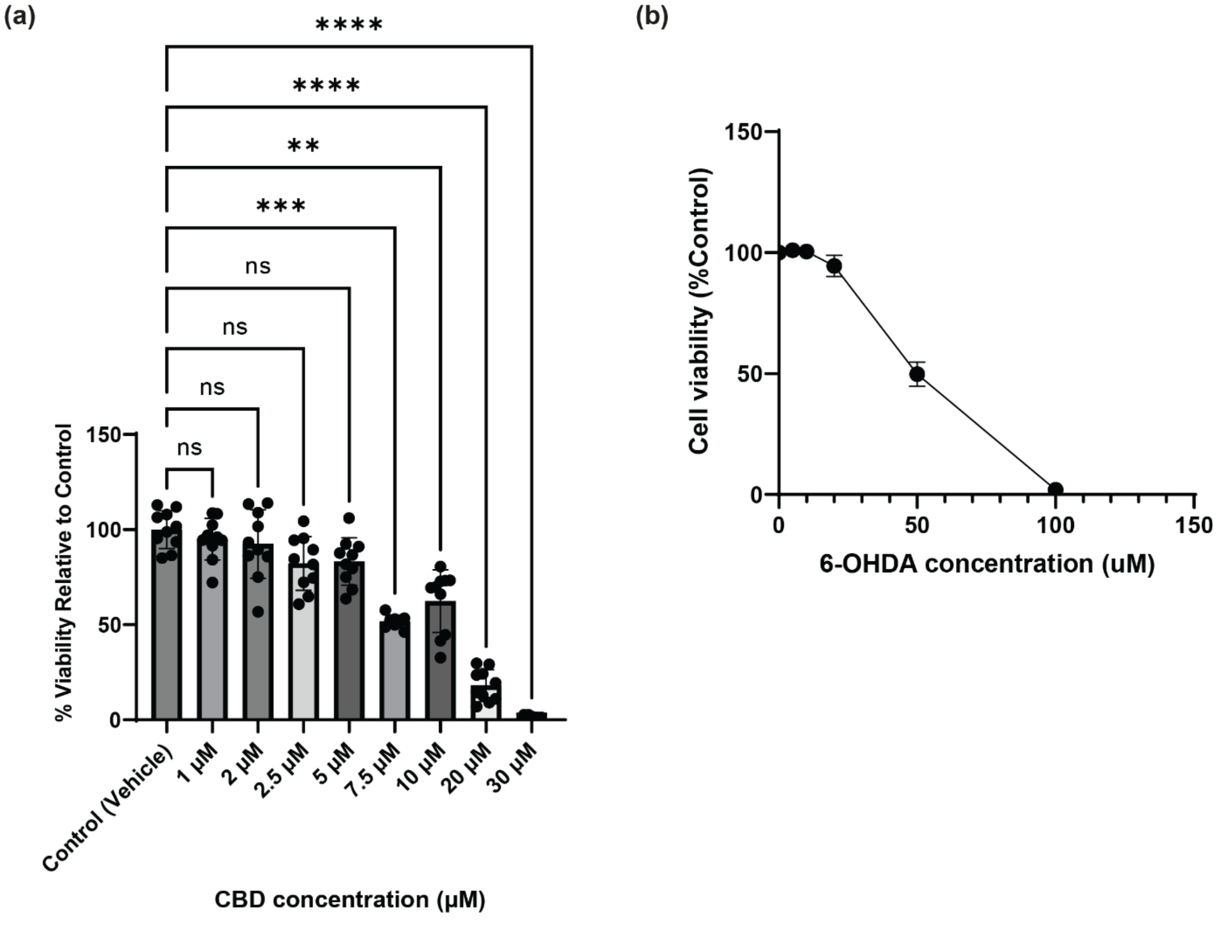


**Supplementary Figure 1** Dose-optimization studies for 6-OHDA and CBD. **(a)** Dose-response curve for calculating the half-maximal inhibitory concentration (IC50) of 6-OHDA. Cell viability of undifferentiated SH-SY5Y cells was measured across a range of 6-OHDA concentrations (0, 5, 10, 20, 50, 100 μM) after 24 hours to determine the optimal cytotoxic dose. A non-linear regression model was fitted to the data, yielding an approximate IC50 of 50 μM, which provided an optimal therapeutic window for evaluating candidate compounds. **(b)**

Assessment of basal cytotoxicity for different CBD concentrations. Undifferentiated SH-SY5Y cells were treated for 24 hours with vehicle control or a range of CBD concentrations (1 µM to 30 µM). Viability is presented relative to the vehicle control, confirming that the 5 µM CBD dose selected for subsequent mechanistic studies is non-toxic. Significant intrinsic cytotoxicity is observed at higher doses (10-30 µM). Data are presented as mean ± SD from independent biological replicates (n = 10), with individual data points shown. Due to non-normal data distribution in some treatment groups in (b), statistical significance was assessed using the Kruskal-Wallis nonparametric test, followed by Dunn's multiple comparisons test against the control. $^{**}p < 0.01$, $^{***}p < 0.001$, $^{****}p < 0.0001$; ns, not significant.

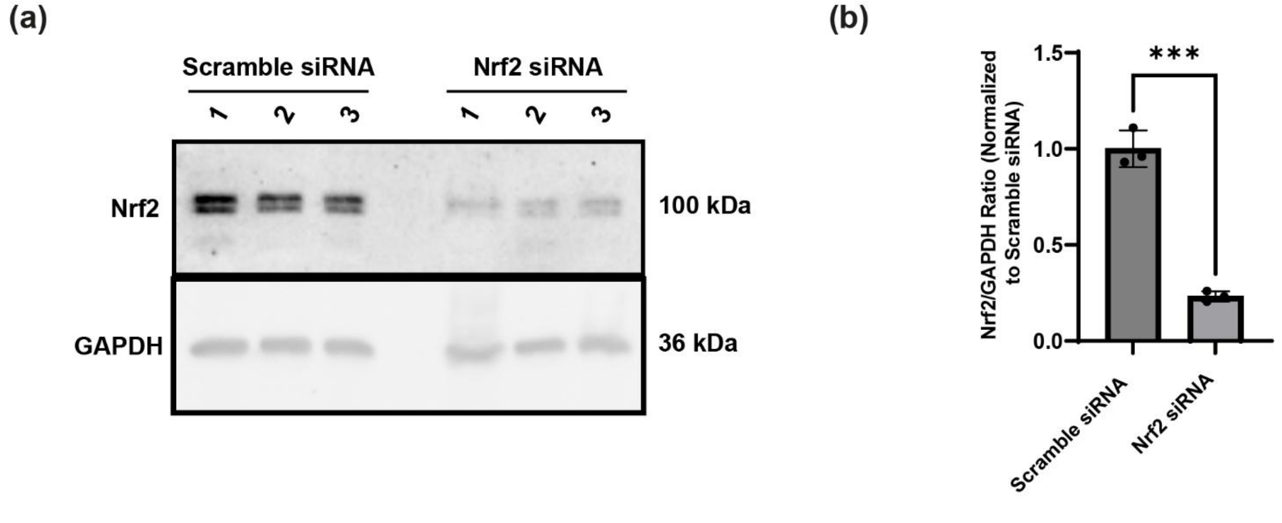


**Supplementary Figure 2** Validation of siRNA-mediated Nrf2 knockdown in SH-SY5Y cells. To confirm the efficiency of targeted genetic silencing, SH-SY5Y cells were transfected with either Scramble (control) or Nrf2-specific siRNA before experimental treatments. **(a)** Representative Western blot images demonstrating the expression of Nrf2 protein in Scramble and Nrf2 siRNA-transfected cells. GAPDH served as the loading control to ensure equal protein loading. **(b)** Densitometric quantification of Nrf2 protein levels. The Nrf2 signal was normalized to the loading control and is expressed relative to the Scramble siRNA group. Data are presented as mean ± SD from independent biological replicates (n = 3). Statistical significance was determined using an unpaired, two-tailed Student's t-test. ***p < 0.001.

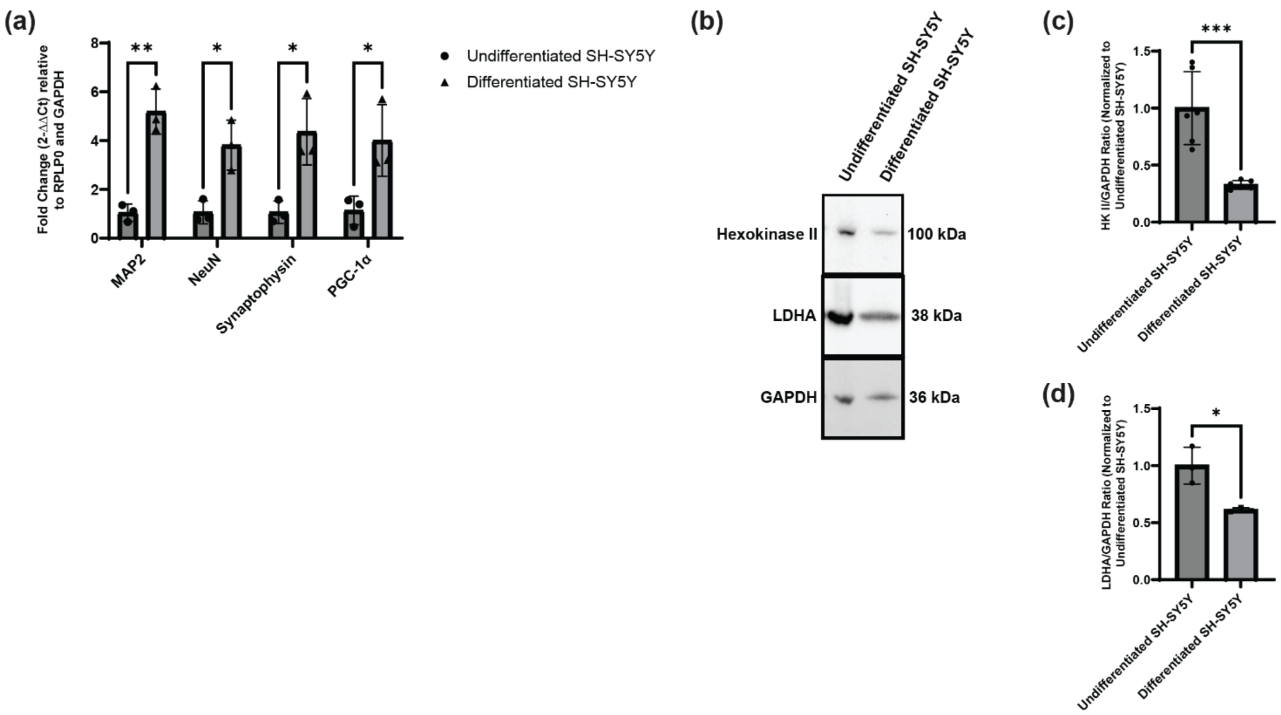


**Supplementary Figure 3** Validation of cellular differentiation and metabolic shift in SH-SY5Y cells. SH-SY5Y cells were differentiated for 14 days using retinoic acid (RA) and brain-derived neurotrophic factor (BDNF) before analysis. **(a)** RT-qPCR analysis of mature neuronal markers (*MAP2*, *NeuN*, *Synaptophysin*) and the mitochondrial biogenesis regulator *PGC-1α*. Gene expression is presented as fold change ($2^{-\Delta\Delta C_t}$) relative to the geometric mean of the housekeeping genes *RPLP0* and *GAPDH*. **(b)** Representative Western blot images demonstrating the downregulation of key glycolytic enzymes, Hexokinase II (HK II) and Lactate Dehydrogenase A (LDHA), following differentiation. **(c–d)** Densitometric quantification of HK II **(c)** and LDHA **(d)** protein levels. Protein expression was normalized to GAPDH and is expressed as a ratio relative to the undifferentiated SH-SY5Y control group. All data are presented as mean ± SD from n = 3 independent biological replicates and n = 6 independent biological replicates in the HK II western blot. Statistical significance for the qPCR data (a) was determined using multiple t-tests corrected for multiple comparisons using the Holm-Sídák method. Statistical significance for the Western blot densitometry (c, d) was determined using an unpaired, two-tailed Student's t-test. *p < 0.05, **p < 0.01, ***p < 0.001.

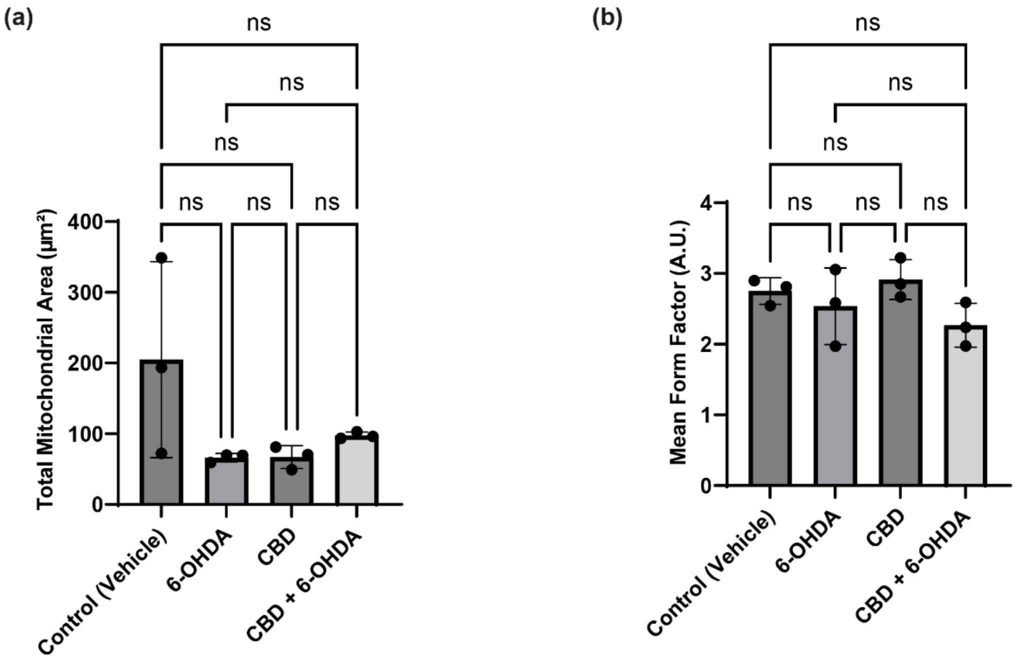


**Supplementary Figure 4** Extended analysis of mitochondrial morphology in undifferentiated and differentiated SH-SY5Y cells. Mitochondrial morphology was visualized using MitoTracker Red CMXRos and subjected to quantitative morphological analysis. **(a)** Quantification of Total Mitochondrial Area (μm²) in undifferentiated SH-SY5Y cells. While 6-OHDA significantly reduced the total mitochondrial area compared to the vehicle control, pretreatment with 5 μM CBD did not significantly restore mitochondrial mass in this proliferative state. **(b)** Quantification of Mean Form Factor (A.U.) in differentiated SH-SY5Y cells. Unlike in undifferentiated cells, 6-OHDA did not significantly alter the mean form factor compared to the vehicle control, and there was no significant difference between the 6-OHDA alone and CBD + 6-OHDA treatment groups in this mature neuron-like model. Data are presented as mean ± SD. Individual data points represent single cells analyzed per condition. Statistical significance was determined using Kruskal-Wallis followed by Dunn's multiple comparisons test. **p < 0.01, ***p < 0.001; ns, not significant.

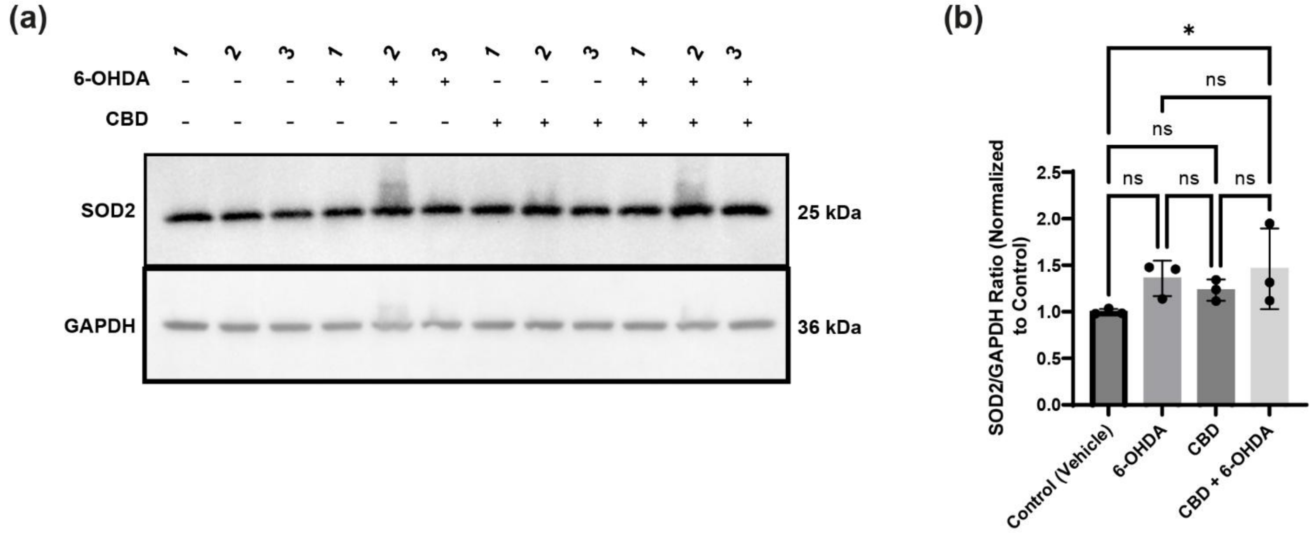


**Supplementary Figure 5** Effect of CBD and 6-OHDA on mitochondrial SOD2 protein expression. **(a)** Representative Western blot of SOD2 and the loading control GAPDH from differentiated SH-SY5Y cell lysates across the four experimental groups: Control (Vehicle), 6-OHDA, CBD, and CBD + 6-OHDA. **(b)** Densitometric quantification of SOD2 protein levels. Target band integrated densities were normalized to their respective GAPDH loading controls and expressed as a fold change relative to the vehicle control group. Data are presented as mean ± SD from three independent biological replicates (n = 3). Statistical significance was determined using one-way ANOVA followed by Tukey's multiple comparisons test. *p < 0.05 vs. Control; ns = not significant.

**Supplementary Table 1.** List of primer sequences used for RT-qPCR analysis.

| **Gene Symbol** | **Gene Name** | **Primer Orientation** | **Sequence (5′→3′)** |
|---|---|---|---|
| **HMOX1** | Heme Oxygenase 1 | Forward | CCAGGCAGAGAATGCTGAGTTC |
| | | Reverse | AAGACTGGGCTCTCCTTGTTGC |
| **NQO1** | NAD(P)H Quinone Dehydrogenase 1 | Forward | CCTGCCATTCTGAAAGGCTGGT |
| | | Reverse | GTGGTGATGGAAAGCACTGCCT |
| **NFE2L2** | Nuclear Factor Erythroid 2-Related Factor 2 (Nrf2) | Forward | CACATCCAGTCAGAAACCAGTGG |
| | | Reverse | GGAATGTCTGCGCCAAAAGCTG |
| **RPLP0** | Ribosomal Protein Lateral Stalk Subunit P0 | Forward | TGGTCATCCAGCAGGTGTTCGA |
| | | Reverse | ACAGACACTGGCAACATTGCGG |
| **GAPDH** | Glyceraldehyde-3-Phosphate Dehydrogenase | Forward | GTCTCCTCTGACTTCAACAGCG |
| | | Reverse | ACCACCCTGTTGCTGTAGCCAA |
